# Manganese toxicity disrupts indole acetic acid homeostasis and suppresses CO_2_ assimilation reaction in rice plants

**DOI:** 10.1101/2021.03.16.435720

**Authors:** Daisuke Takagi, Keiki Ishiyama, Mao Suganami, Tomokazu Ushijima, Takeshi Fujii, Youshi Tazoe, Michio Kawasaki, Ko Noguchi, Amane Makino

**Author notes:** Corresponding author Daisuke Takagi. Faculty of Agro-Food Science, Niigata Agro-Food University, Tainai, Niigata, 959-2702 Japan (Youshi Tazoe). Faculty of Food and Agricultural Sciences, Fukushima University, Kanayagawa, Fukushima, 960-1296, Japan (Mao Suganami). **E-MAIL ADDRESSES** Keiki Ishiyama; Mao Suganami; Tomokazu Ushijima; Takeshi Fujii; Youshi Tazoe; Michio Kawasaki; Ko Noguchi; Amane Makino.

## Abstract

Despite the essentiality of Mn in terrestrial plants, its excessive accumulation in plant tissues causes growth defects, known as Mn toxicity. Mn toxicity can be divided into apoplastic and symplastic types depending on its onset. For growth defects, symplastic rather than apoplastic Mn toxicity is hypothesised to be more critical. However, details of the relationship between growth defects and symplastic Mn toxicity remains elusive. In this study, we aimed to elucidate the molecular mechanisms of symplastic Mn toxicity in rice plants. We found that under excess Mn conditions, CO_2_ assimilation was inhibited by stomatal closure, and both carbon anabolic and catabolic activities were decreased. In addition to stomatal dysfunction, stomatal and leaf anatomical development were also altered by excess Mn accumulation. Furthermore, the indole acetic acid (IAA) concentration was decreased, and auxin-responsive gene expression analyses showed IAA-deficient symptoms in leaves due to excess Mn accumulation. These results suggest that excessive Mn accumulation causes IAA deficiency, and low IAA concentrations suppress plant growth by suppressing stomatal opening and leaf anatomical development for efficient CO_2_ assimilation in leaves.

**HIGHLIGHT:** Increased Mn concentration lowers auxin concentrations in rice leaves, which suppresses photosynthesis by changing stomatal function and development.

## INTRODUCTION

Mn is an essential nutrient for both terrestrial plants and animals (Shao *et al*., 2017). With regard to terrestrial plants, Mn was first discovered in their ash, and McHargue (1922) proved that Mn is an essential nutrient. Manganese possesses a wide variety of physiological functions in plant cells. For example, Mn activates more than 35 different enzymes such as chloroplast RNA-polymerase, and several enzymes involved in the tricarboxylic acid (TCA) cycle and shikimic acid pathway (Broadley *et al*., 2012). Mn is also directly involved in the physiological function of Mn-superoxide dismutase (SOD) as a cofactor to detoxify reactive oxygen species (ROS) and oxalate oxidase (Broadley *et al*., 2012). Among these physiological functions, the Mn cluster within the oxygen-evolving complex of photosystem II (PSII) is crucial for driving photosynthesis in terrestrial plants (Suga *et al*., 2015).

Although Mn is indispensable for terrestrial plants, excess Mn accumulation in leaves reduces their growth and crop yield, through what is known as Mn toxicity (Foy *et al*., 1978; Li *et al*., 2019). The toxic effects of Mn are observed in various terrestrial plants, but the critical concentration for expressing toxicity varies depending on plant species and genotype (Foy *et al*., 1978; Horst *et al*., 1999; Horiguchi, 1987; Millaleo *et al*., 2010). For example, *Zea mays* L. shows signs of Mn toxicity at an accumulation of 200 µg Mn g^-1^ (dry weight; D.W.) in the leaves; in contrast, *Lupinus albus* L. or woody Mn hyperaccumulator species, such as *Gossia bidwillii*, can accumulate more than 10,000 µg Mn g^-1^ (D.W.) without Mn toxicity symptoms (Blamey *et al*., 1986, 2015; Fernando *et al*., 2006). These differences in the critical concentration for Mn toxicity are derived from the different capacities of Mn compartments within cells (Blamey *et al*., 2015). Excessive Mn is sequestrated to vacuoles in a chelated form with organic acids (malate/citrate) to maintain the Mn concentration within cells (Blamey *et al*., 2015). However, when the vacuole capacity for storing Mn reaches the upper limit, the Mn concentration increases in the symplastic and apoplastic regions, where Mn toxicity occurs (Horiguchi, 1987; Führs *et al*., 2009). In fact, Mn-sensitive plants show a smaller vacuole capacity for retaining Mn than do Mn-tolerant plants (Horst *et al*., 1999; Blamey *et al*., 2015). This interpretation has been validated by loss-of-function and overexpression mutants of tonoplast-localised Mn transporter proteins such as metal tolerant protein (MTP) 8 and calcium exchanger (CAX) 2, which show a susceptible and tolerant phenotype to Mn toxicity, respectively (Hirschi *et al*., 2000; Delhaize *et al*., 2003; Chen *et al*., 2013; Eroglu *et al*., 2016). In addition to these mechanisms, Mn-tolerant plants, such as *Helianthus annuus* L., use trichomes as Mn storage tissue to avoid an excessive increase in Mn concentration within cells (Blamey *et al*., 1986, 2015). However, this mechanism depends on the plant species and is not a general strategy to prevent Mn toxicity (Blamey *et al.,* 2015).

Mn toxicity can be divided into two categories depending on the cell part of its onset: apoplastic and symplastic (Fernando *et al*., 2016). The primary symptoms of apoplastic Mn toxicity in terrestrial plants are brown spots containing oxidised phenolics, oxidised Mn [Mn^3+^ and Mn^4+^], and callose in the leaves (Kenten and Mann, 1957; Horiguchi, 1987; Wissemeir and Horst, 1987, 1992; Horst *et al*., 1999; Fecht-Christoffers *et al*., 2003*a*, *b*; Blamey *et al*., 2015). Mn accumulation in the apoplast stimulates the expression of class III peroxidases (PODs), which undertake both H_2_O_2_ production through NADH-peroxidase activity and H_2_O_2_ consumption through guaiacol-peroxidase activity (Fecht-Christoffers *et al.,* 2003a, b; Führs *et al.,* 2009). Furthermore, the concentrations of phenolics, which suppress the NADH-peroxidase activity, are lowered in the apoplast and, as a consequence, H_2_O_2_ production is accelerated in the apoplast (Fecht-Christoffers *et al.,* 2006; Führs *et al.,* 2009). The H_2_O_2_ consumption reaction by the guaiacol activity of PODs also proceeds by utilising phenolics in the apoplast; subsequently, the intermediate phenolic oxidation product, phenoxyl radicals (PhO •), oxidises Mn^2+^ to Mn^3+^ (Takahama, 2004). Because PhO • is regenerated to phenolics after the reaction with Mn^2+^, the continuous reactions of H_2_O_2_-production/consumption by the class III PODs accumulate the oxidised phenolics and oxidised Mn in the apoplast (Kenten and Mann, 1957).

Although brownish spots are an indicator of Mn toxicity in terrestrial plants, attenuation of plant growth by Mn toxicity is not always accompanied by the expression of brown spots in leaves (Blamey *et al.,*1986, 2015; Horiguchi, 1987; Wissemeier and Horst, 1992; González *et al*., 1998; Chen *et al*., 2013; Tsunemitsu *et al*., 2018). Under excess Mn accumulation in the symplast, leaves show lower photosynthetic activity and lower Chl content (Nable *et al*., 1988; Macfie and Taylor, 1992; González and Lynch, 1997; Kitao *et al*., 1997; Fernando and Lynch, 2015). Houtz *et al*. (1988) demonstrated a decrease in photosynthetic activity despite the absence of brown necrotic spots on leaves grown under high Mn concentrations, implying that a decrease in photosynthetic activity occurred independently of apoplastic Mn toxicity. These characteristics are termed symplastic Mn toxicity (Fernando *et al*., 2016). The decreases in Chl concentration and photosynthetic activities have been hypothesised to be caused by photoinhibition by ROS; the decrease in photosystem I (PSI) content or the disturbance of Chl synthesis due to the inhibition of Fe absorption; Mn binding to Rubisco instead of Mg; or stomatal dysfunction (Clairmont *et al*., 1986; Suresh *et al*., 1987; Houtz *et al*., 1988; McDaniel and Toman, 1994; Millaleo *et al*., 2013). However, these hypotheses are often independently discussed at both *in vivo* and *in vitro* levels; therefore, the mechanisms by which symplastic Mn toxicity suppresses photosynthesis are less evident. Elucidating Mn toxicity including both its apoplastic and symplastic mechanisms would contribute to establishing a strategy for protecting terrestrial plants against Mn toxicity in agricultural land (Fernando and Lynch, 2015). Currently, the strategy for preventing Mn toxicity depends on two mechanisms. One is limiting Mn absorption and transport, and the other is Mn sequestration from the cytosol to the vacuole (Li *et al*., 2019). These strategies are important in preventing Mn toxicity. However, identifying the expression mechanisms of symplastic Mn toxicity would be informative for manipulating Mn toxicity in terrestrial plants without limiting the Mn transportation system. This knowledge would be applicable to plant engineering processes such as Mn phytoremediation (Lei *et al*., 2007).

In this study, we investigated the mechanisms of symplastic Mn toxicity to suppress photosynthetic activity in rice leaves. Rice shows high tolerance toward Mn toxicity, which make it easy to investigate the effect of excess Mn concentrations without severe necrosis and chlorosis (Führs *et al*., 2010). Additionally, complete genetic information is available for investigating gene expression. Previous studies have indicated that photosynthetic electron transport activities on the thylakoid membrane and Rubisco isolated from leaves grown under high Mn concentrations were robust, although CO_2_ assimilation was substantially suppressed in leaves (Houtz *et al*., 1988; Nable *et al*., 1988; Chatterjee *et al*., 1994). Based on these observations, we hypothesised that chloroplast proteins are not the primary targets of Mn toxicity, but factors that modulate CO_2_ assimilation are targeted under Mn toxicity. Here, we discuss how photosynthetic activity is limited under Mn toxicity conditions from a wide range of physiological responses in plant cells.

## MATERIALS AND METHODS

### Plant materials and plant growth conditions

*Oryza sativa* L. ‘Nipponbare’ wild-type (WT) plants were grown in hydroponic culture, as reported in a previous study (Makino *et al.,* 1994). We employed two Mn concentration conditions by changing the MnSO_4_ application to 0.5 µM (control conditions) and 200 µM (Mn-toxic conditions) (Tsunemitsu *et al.,* 2018). The pH of the hydroponic culture was adjusted to 5.2 using HCl, and the solution was replenished twice a week. The chamber was maintained at 60% relative humidity with a 14 h light (28 °C) and 10 h dark (25 °C) photoperiod. The light intensity was 500–600 µmol photons m^−2^ s^−1^. All physiological, structural, and genetic analyses were performed for fully and newly expanded leaves after 70 d of germination, and the analyses were completed before heading.

### Leaf photosynthesis and respiration measurements in rice plants

The gas exchange analysis, Chl fluorescence, and oxidised reaction centre Chl in PSI (P700^+^) were simultaneously measured using a combined system of Li-6400 (Li-COR Inc., Lincoln, USA), Mini-PAM, and PAM-101 equipped with a dual-wavelength emitter-detector unit (ED-P700DW) (Heinz Walz GmbH, Effeltrich, Germany) with a cold halogen lamp. Ambient air (40 Pa CO_2_ and 21 kPa O_2_) and pure CO_2_ gas were mixed to maintain the CO_2_ concentration during measurements. The gases were saturated with water vapour at 18.0 ± 0.1 °C, and the leaf temperature was maintained at 28 °C. The Chl fluorescence parameters Y(II), Y(NPQ), and Y(NO) were calculated as described by Baker (2008), and a measuring light (0.1 µmol photons m^−2^ s^−1^) and a saturated pulse (10,000 µmol photons m^−2^ s^−1^, 600 ms) were employed to calculate the photosynthetic parameters of PSII. The oxidation-reduction state of P700 in PSI was determined according to the method described by Klughammer and Schreiber (1994). The maximum oxidation level of P700 (Pm) was obtained using a saturated pulse under far-red light illumination, and PSI photosynthetic parameters [Y(I), Y(ND), and Y(NA)] were calculated using the oxidation-reduction state of P700 as determined by a saturated pulse method.

The CO_2_ emission rate was measured using a Li-6400 in the gaseous phase. The O_2_ absorption rate was measured using an O_2_-electrode (CB1D; Hansatech Instruments Ltd., King’s Lynn, UK) in the aqueous phase at 25 °C (Hachiya *et al.,* 2010). For efficient absorption of the chemical reagent into leaves in the aqueous phase, the leaf surface was washed with 10% (v/v) dimethyl sulfoxide, and the leaves were finely cut up. The O_2_ absorption rate was measured in the reaction buffer [50 mM *N*-(2-hydroxyethyl)piperazine-*N*’-2-ethanesulfonic acid (HEPES)-KOH (pH 6.6), 10 mM 2-(*N*-morpholino)ethanesulfonic acid, 0.2 mM CaCl_2_, 50 mM sucrose]. To evaluate the uncoupled respiration rate and alternative oxidase (AOX) and cytochrome *c* oxidase (COX) activities in the mitochondria, 10 µM carbonyl cyanide m-chlorophenyl hydrazine, 2 mM n-propyl gallate, and 2 mM KCN were added sequentially.

### Quantification of leaf mineral elements and chlorophyll content

Leaf mineral elements were analysed according to Takagi *et al*. (2020). Briefly, the leaf blades were dried at 70 °C and ground using a homogeniser. To determine the mineral content, except for that of N, the dry matter was digested overnight in an acid mixture (HNO_3_:H_2_SO_4_:HClO_4_ = 5:1:2). Subsequently, the solution was heated at 150 °C for 30 min in a heat block. After cooling, the solutions were heated again at 200 °C for 1 h. After decomposition, the mineral contents were quantified by inductively coupled plasma optical emission spectroscopy (ICP-OES) (iCAP™ 7200; Thermo Fisher Scientific Inc., Waltham, USA). To measure the total N concentration in the leaves, dried leaves were decomposed using 60% (v/v) H_2_SO_4_ with 30% (v/v) H_2_O_2_. The N concentration was determined using Nessler’s reagent in a decomposed solution after the addition of 10% (w/v) potassium sodium tartrate and 2.5 M NaOH, and the absorbance was measured at 420 nm (Takagi *et al.,* 2017). The leaf Chl content was quantified using fresh samples with *N*, *N*-dimethylformamide, as described previously (Porra *et al.,* 1989).

### Carbohydrates quantification in leaves

Leaves were sampled at the end of the day (1 h before the start of the dark period) and at the end of the night (1 h before the start of the light period); subsequently, leaves were frozen using liquid nitrogen. Glucose, sucrose, and starch were extracted from leaves according to Morita *et al*. (2015) and quantified using the Enzytech^TM^ D-Glucose/Sucrose kit (R-Biopharm AG, Darmstadt, Germany).

### Quantification of leaf amino acid contents

The leaves were homogenised using 10 mM HCl with liquid nitrogen. After centrifugation (15,000 × *g*, 4 °C, 5 min), the supernatant was applied to a centrifugation filter (Amicon Ultra 0.5 mL 3 K device; Merck KGaA, Darmstadt, Germany), and the fraction containing free amino acids was obtained by centrifugation (15,000 × *g*, 4 °C, 10 min). The amino acid was labelled using AccQ-Tag Ultra Derivatization Kit (Waters Co., Milford, USA) according to the manufacture’s instructions, and the free amino acids were separated using an HPLC system. The HPLC system consisted of a 305 Piston Pump as the system controller and a 306 Pump as a multi-pump application equipped with an 811D Dynamic Mixer and 805 Manometric Module, using the Trilution LC software (Gilson, Villiers le Bel, France). The sample solution was manually injected into the HPLC system. The derivative amino acids were separated on a Sepax Bio-C18 (φ4.6 × 250 mm) (Sepax Technologies, Inc., Newark, USA) and eluted by 100% solvent A [5% (v/v) methanol in 20 mM CH_3_COONa (pH 6.5)] for 4 min after injection and by a linear gradient of 94% solvent A to 30% solvent B (100% acetonitrile) for another 65 min. The column was washed with 100 % solvent B for 20 min and equilibrated with 100% solvent A for 16 min. The flow rate was 0.6 mL min^-1^, and the column temperature was maintained at 25 °C by incubation in a hot pocket (Thermo Fisher Scientific Inc., Waltham, USA). The peaks derived from derivatised amino acids were detected using UV/VIS-155 (Gilson, Villiers le Bel, France).

### TCA cycle enzyme activities

The TCA cycle enzyme activities in the leaf blades were determined according to Noguchi and Terashima (2006). Briefly, the leaves were homogenised with the extraction buffer [100 mM HEPES-KOH (pH 7.5), 10 mM KH_2_PO_4_, 0.5 mM EDTA-Na, 10 mM dithiothreitol, 0.05% (v/v) TritonX-100, and 20% (v/v) glycerol] with polyvinylpyrrolidone. The reaction medium was prepared for each enzyme according to the previous study, and the enzyme activity was determined at 30 °C using UV-160A equipped with a temperature control system (Shimadzu, Kyoto, Japan).

### Structural analysis of stomata complex and leaf cross-section

For scanning electron microscopic analysis, the leaves were fixed with half-strength Karnovsky’s solution [50 mM phosphate buffer (pH 7.2), 2.5% (v/v) glutaraldehyde, 2% (v/v) paraformaldehyde] with vacuum infiltration for 3 h. After fixation, the leaves were washed with the phosphate buffer and dehydrated using a dilution series of ethanol, followed by t-butanol. Leaves were dried using a t-butanol freeze drier (Vacuum Device Inc., Ibaraki, Japan) and coated with Pt (JEC-300FC; JEOL Ltd., Tokyo, Japan). The stomatal complex structure was monitored using a JSM-IT200 scanning electron microscope (JEOL Ltd., Tokyo, Japan). The size of the stomatal complex, comprising the paired guard cells, the paired subsidiary cells, and the pore itself, and the stomatal density were analysed using SEM Operation software (JEOL Ltd., Tokyo, Japan). Under each growth condition, the size of the stomata complex and the stomatal density were analysed on an independent section of the leaves as indicated in figure legends of six biologically independent plants. To analyse the vertical and horizontal leaf sections, leaves were sectioned and fixed with formalin-acetate-alcohol (FAA) [63% (v/v) ethanol, 5% (v/v) acetic acid, 5% (v/v) formalin] with vacuum infiltration for 1 h, followed by incubation at 4 °C for 2 d. The leaves were dehydrated using a dilution series of ethanol, followed by n-butanol. The dehydrated leaves were embedded in paraffin and were sectioned at 10 µm thickness using an RX-860 microtome (Yamato Kohki Industrial Co., Ltd., Saitama. Japan). The FAA-fixed leaf sections were stained with 0.05% (w/v) toluidine blue O. To investigate the intact leaf sections, leaves were embedded in 50 mM phosphate buffer (pH 7.2) containing 5% (w/v) agar. The agar-embedded leaves were sectioned at a thickness of 30 µm using DTL-1000 vibratome (Dosaka EM Co., Ltd., Kyoto, Japan). Intact leaf sections and FAA-fixed leaf sections were analysed using an optical microscope (BX53; Olympus Co., Japan).

### Indole acetic acid quantification

The IAA content in leaves was quantified using a GC-MS system, according to Nghi *et al*. (2021). Briefly, the leaves were homogenised in 70% (v/v) cold acetone. After centrifugation, the supernatant was reduced to the aqueous phase by vacuum centrifugation equipped with a cold trap. Subsequently, HCl (0.1 mM) was added to adjust the pH to 2.8, and diethyl ether was added to dissolve IAA into the diethyl ether fraction. The diethyl ether fraction was dried under vacuum and subsequently dissolved in *N*,*O*-bis(trimethylsilyl) tri-fluoroacetamide (BSFTA) containing 1% trimethylchlorosilane. IAA was silylated at 70 °C for 1 h. The silylated IAA was identified using a GC-MS-QP2010 SE (Shimadzu, Kyoto, Japan) with a DB-WAX column. The conditions for GC-MS analysis were the same as those described previously (Fujii *et al*., 2010). The silylated IAA was identified by fragment ion 202 (*m/z*) and molecular ion 319 (*m/z*) (Ljung *et al*., 2005; Gutierrez *et al*., 2009; Nghi *et al*., 2021).

### Gene expression analysis

To investigate gene expression in rice leaves, mRNA was isolated from the leaf blades according to Suzuki *et al*. (2004). After mRNA isolation, cDNA was synthesised using PrimeScript™ RT Reagent Kit (Takara Bio Inc., Shiga, Japan), and the mRNA content was quantified by real-time PCR using the LightCycler® 96 System (Roche Diagnostics K.K., Tokyo, Japan) with a KAPA SYBR FAST One-Step qRT-PCR Kit (Nippon Genetics Co., Ltd, Tokyo, Japan). The primer sequences used for real-time PCR analyses are listed in Supplementary Table S1.

### Simulation of CO_2_ fixation rate according to biochemical photosynthetic model

Rubisco limited CO_2_ fixation rate was calculated from the equation of von Caemmerer and Farquhar (1981):

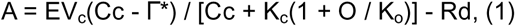

Where E is the amount of Rubisco protein (assumed to be 5.3 μmol m^-2^, from rice grown with hydroponic solution same as in the present study described by Suganami et al. 2021), V_c_ is the Rubisco activity of carboxylation, Cc is the chloroplastic CO_2_ partial pressure, O is the partial pressure of O_2_in the chloroplast (assumed to be the same as in the atmosphere, 21 kPa), K_c_ and K_o_ are the Michaelis-Menten constants for CO_2_ and O_2_, and Rd is the day respiration (assumed to be 1.0 μmol m^-2^ s^-1^). Γ* is the CO_2_ compensation point of photosynthesis in the absence of Rd as defined as follows:

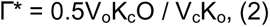

where V_o_ is the Rubisco activity of oxygenation. Mg^2+^-binding Rubisco kinetics were taken from Makino et al., (1988). Mn^2+^-binding Rubisco kinetics were calculated by multiplying the average ratio of Mn^2+^/Mg^2+^-binding Rubisco kinetics and Mg^2+^-binding Rubisco kinetics. The average ratio of Mg^2+^/Mn^2+^-binding Rubisco kinetics were calculated by the data from Bloom and Kameritsch (2017). Rubisco kinetics were summarized in Supplementary Table S2. Photorespiration rate were was calculated from the equation of von Caemmerer and Farquhar (1981):

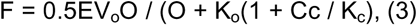

### Statistical analysis

All measurement data are expressed as means ± standard deviation (SD), or box plots with bars indicating the 1.5 × interquartile range (IQR) and with squares indicating the mean value of at least three independent biological analyses. Significant differences in physiological and biological parameters between the control- and Mn-toxic conditions were detected by the non-parametric Kruskal-Wallis test. All statistical analyses were performed using Origin Pro 2020 (LightStone Corp., Tokyo, Japan).

## RESULTS

### Phenotypes of rice grown under Mn-toxic conditions

To study the effects of Mn toxicity, *Oryza sativa* L. ‘Nipponbare’ was grown under high Mn concentrations (200 µM; Mn-toxic conditions) in hydroponic culture for 70 d after germination. Under Mn-toxic conditions, the leaves showed pale green and brown-coloured sections, especially at the tips (Fig. 1A, B). These observations correspond with typical Mn toxicity symptoms in rice leaves (Vlamis and Williams, 1964; Chen *et al*., 2013). Compared with the control conditions, Mn-toxic conditions decreased the total dry weight of rice, especially the dry matter of the leaf sheath and root (Fig. 1C, D).

**Figure 1.**
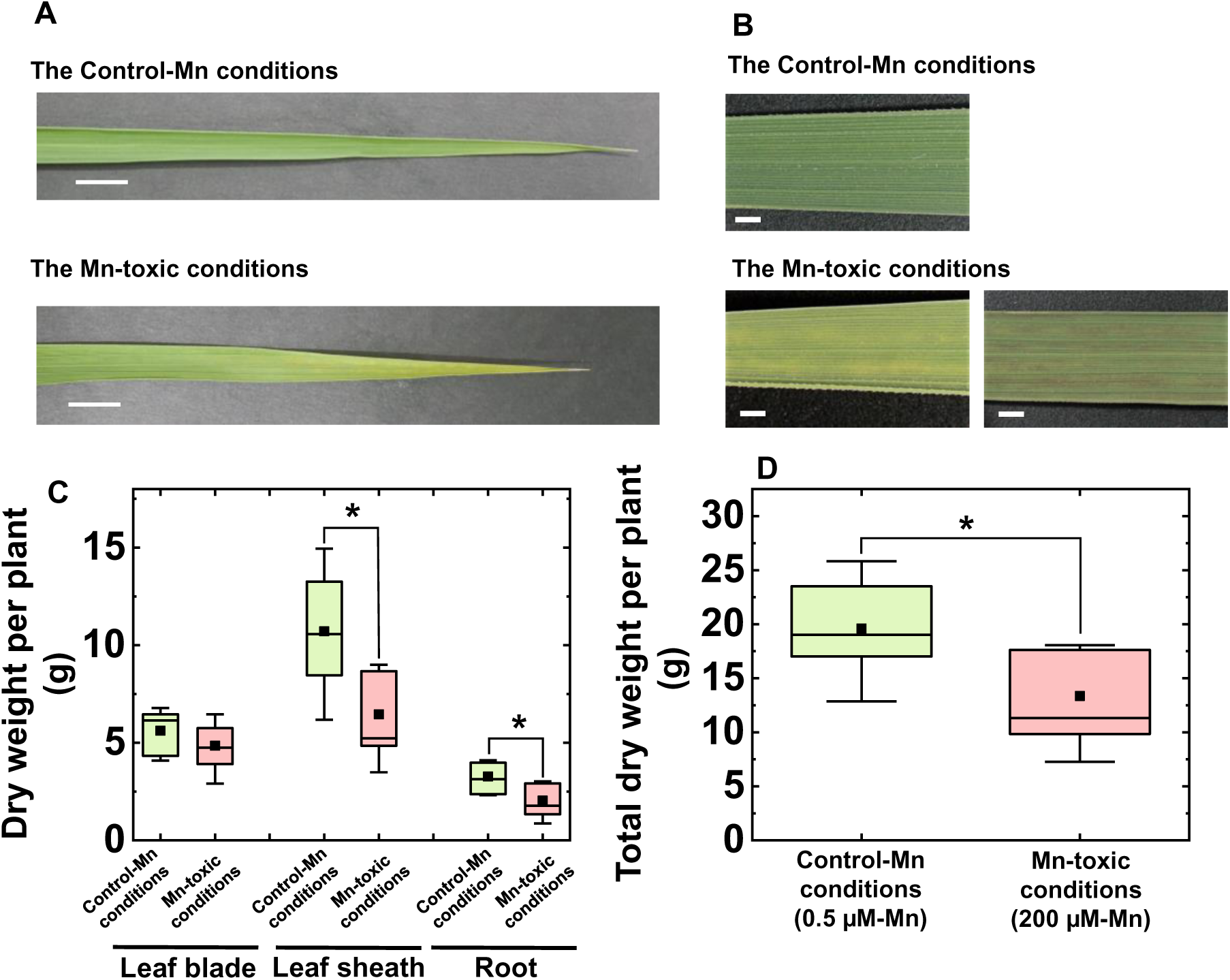
Phenotypes of rice plants grown under control and Mn-toxic conditions. The pictures of fully expanded leaf blades grown under control and Mn-toxic conditions are shown in (A), and (B) shows an enlarged picture of these. The white bars indicate lengths of 1 cm (A) and 1 mm (B), respectively. (C) The dried weights of the leaf blade, leaf sheath, and root, respectively. The sum of the dried weights shown in (C). Data are shown as box plots (n = 9), black squares indicate the mean value, and bars indicate the range of the maximum or minimum data within a 1.5× interquartile range (IQR). Green boxes indicate the results of the control conditions, and red boxes indicate those of the Mn-toxic conditions. Asterisks show significant differences between the control and Mn-toxic conditions (*: *p* < 0.05, Kruskal– Wallis test).

To study the effects of high Mn application on leaf mineral composition, the leaf mineral concentrations were quantified. The leaf Mn concentration was significantly increased under Mn-toxic conditions (Fig. 2). This concentration was comparable to that of WT rice, which showed growth inhibition when grown at a concentration of 1,000 µM Mn for 3 weeks (Sasaki *et al*., 2011). The K, Ca, and Fe concentrations were similar under the control and Mn-toxic conditions (Fig. 2). In contrast, the Mg, Zn, and Cu concentrations increased under Mn-toxic conditions (Fig. 2). These results demonstrated decreased growth and the Mn toxicity symptoms accompanied by Mn accumulation in leaves grown under Mn-toxic conditions.

**Figure 2.**
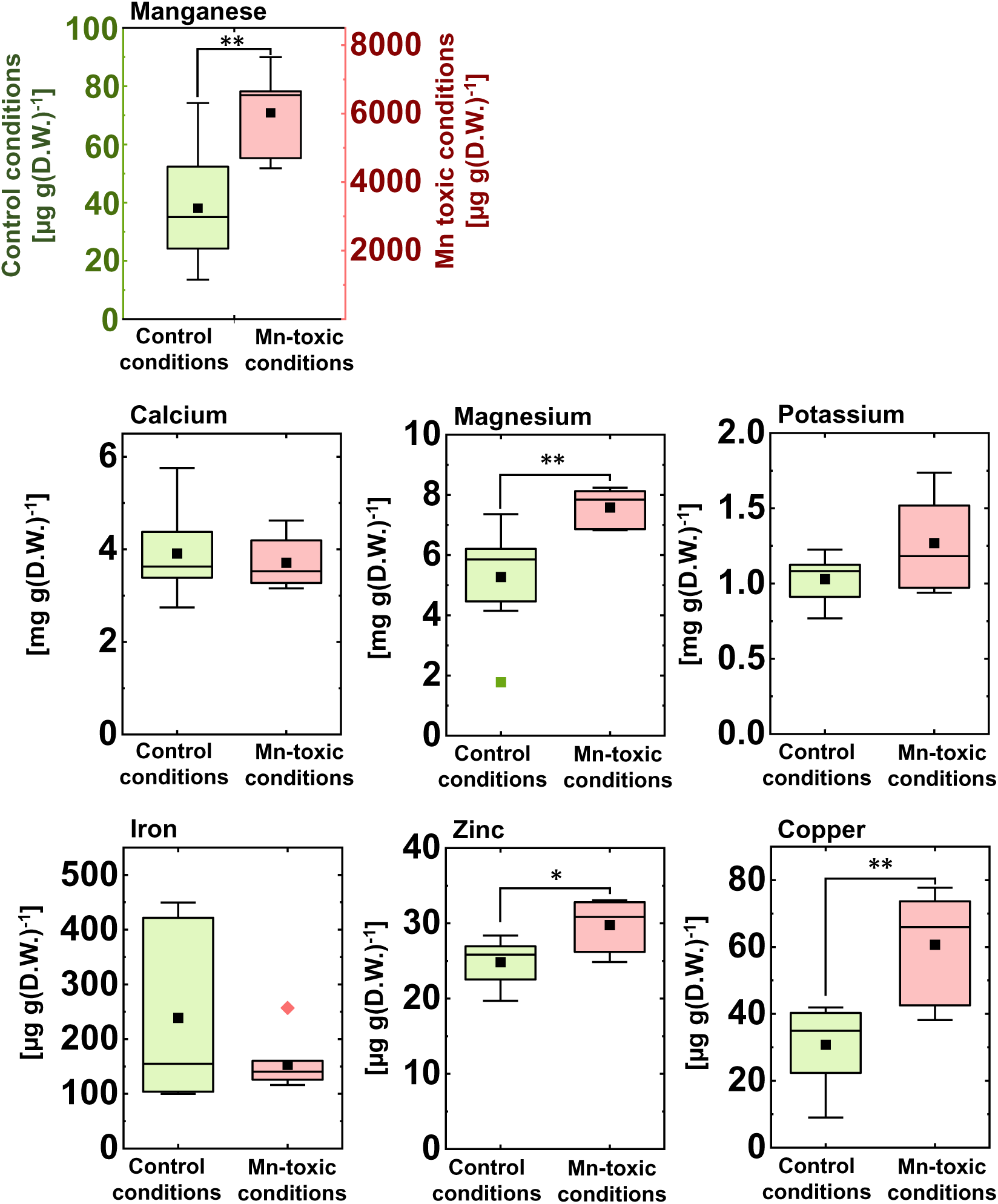
Mineral concentration in the rice leaf blade. Data are shown as box plots (control conditions n = 8, Mn-toxic conditions n = 7), black squares indicate the mean value, and bars indicate the range of the maximum or minimum data within a 1.5× interquartile range (IQR). Green boxes indicate the results of the control conditions, and red boxes indicate those of the Mn-toxic conditions. Asterisks show significant differences between the control and Mn-toxic conditions (*: *p* < 0.05, **: *p* < 0.01, Kruskal–Wallis test).

### Mn-toxic conditions suppressed CO_2_ assimilation by limiting stomatal conductance in leaves

Next, the steady-state photosynthetic activities were measured under ambient air conditions (40 Pa CO_2_ and 21 kPa O_2_). The CO_2_ assimilation rate was significantly decreased under the high light irradiance by the Mn-toxic treatments (Fig. 3A). The Mn-toxic treatments showed lower stomatal conductance (*g_s_*) than did the control plants, and the change in *g*_s_ with increasing light intensity was also suppressed (Fig. 3B). These stomatal responses to the increase in Mn concentration were consistent with previous studies (Suresh *et al*., 1987; Nable *et al*., 1988; González and Lynch, 1997; Lidon *et al*., 2004; Santos *et al*., 2017). The internal CO_2_ concentration in leaves (Ci) was lower under the Mn-toxic conditions than under control conditions (Fig. 3C). These results indicated that, under Mn-toxic conditions, the stomata limit CO_2_ diffusion into leaves and CO_2_ assimilation is suppressed, particularly under high light conditions.

**Figure 3.**
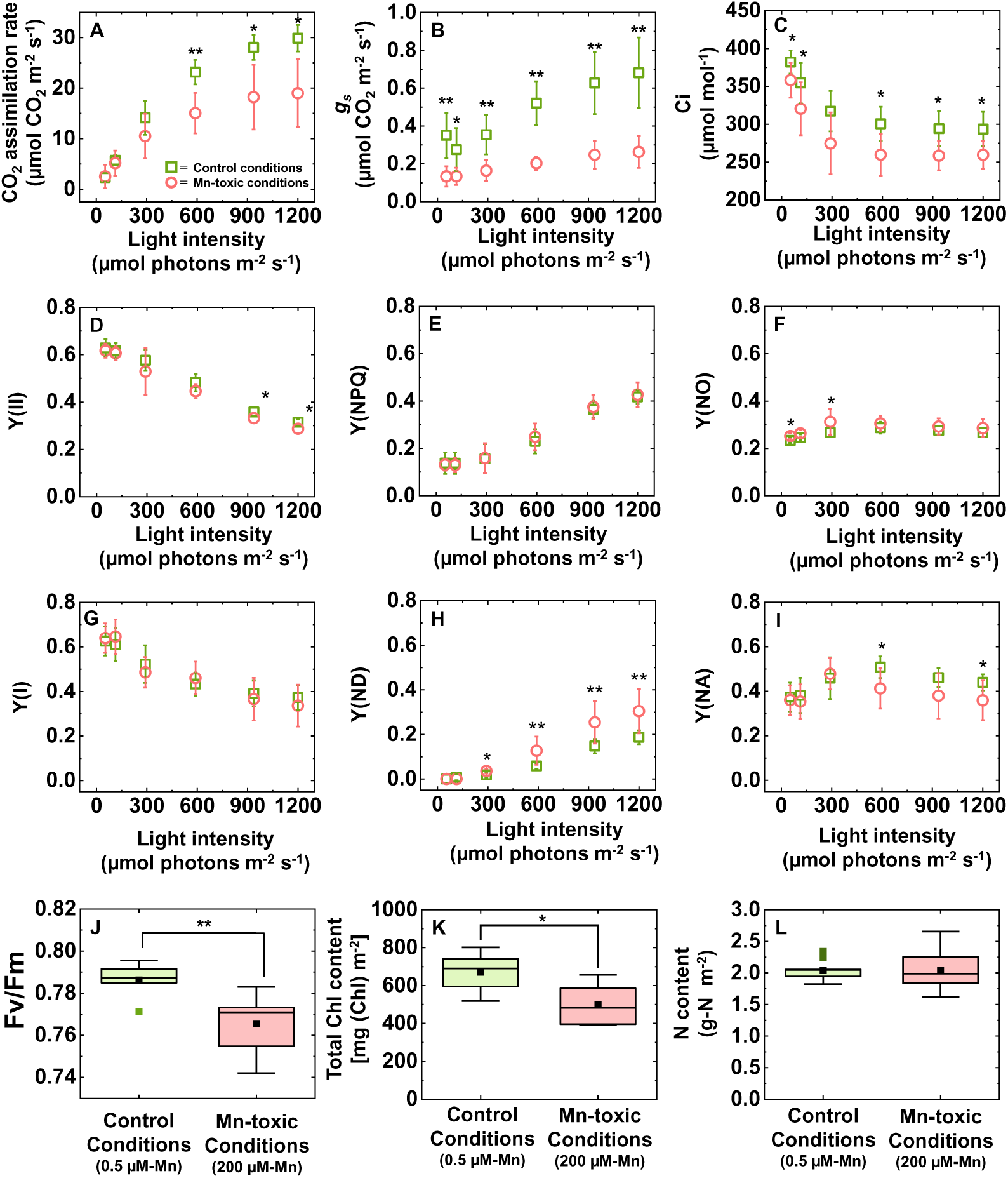
Photosynthetic activities in leaves grown under control and the Mn-toxic conditions. The CO_2_ assimilation rate (A), stomatal conductance (*g_s_*) (B), and intercellular CO_2_ concentration (C) are shown. For the PSII photosynthetic parameters, the quantum yields of PSII [Y(II)] (D), non-photochemical quenching [Y(NPQ)] (E), and non-radiative decay [Y(NO)] (F) are shown. The maximum quantum yield of PSII (Fv/Fm) is shown (J). For PSI photosynthetic parameters, the quantum yields of PSI [Y(I)] (G), non-photochemical quenching at the donor side [Y(ND)] (H), and non-photochemical quenching at the acceptor side [Y(NA)] (I) are shown. (K) and (L) show the total Chl and N concentrations in the leaves, respectively. Data of photosynthetic parameters are shown as means with standard deviation (SD) (n = 7). The results for Fv/Fm (n = 7) and total Chl (n = 7) and N (n = 9) concentrations are shown as box plots. Black squares indicate the mean value, and bars indicate the range of the maximum or minimum data within a 1.5× interquartile range (IQR). Asterisks show significant differences between the control and Mn-toxic conditions (*: *p* < 0.05, **: *p* < 0.01, Kruskal– Wallis test).

In addition to the CO_2_ assimilation rate, the photosynthetic electron transport activities in both PSII and PSI were analysed by chlorophyll fluorescence and P700^+^-dependent absorbance changes in the leaves. The quantum yield of PSII [Y(II)] was lower under Mn-toxic conditions than under control conditions at the high light irradiance (Fig. 3D). This result indicated that the electron transport activity in PSII was suppressed at the high light irradiance, but compared with the CO_2_ assimilation rate, the change in Y(II) was marginal. The quantum yield of non-photochemical quenching [Y(NPQ)] increased similarly with the increase in light intensity under both control and Mn-toxic conditions (Fig. 3E). The quantum yield of non-radiative energy loss [Y(NO)] was different between the control and Mn-toxic conditions under low light irradiance; however, this difference was masked under high light irradiance (Fig. 3F). These results showed that the photoprotective mechanisms in PSII are robust, but the redox state in PSII can be perturbed under Mn toxicity. The quantum yield of PSI [Y(I)] showed similar kinetics to that of Y(II), and no significant differences were observed between the control and Mn toxic condition (Fig. 3G). The quantum yield of non-photochemical quenching at the donor side of PSI [Y(ND)] was higher under Mn-toxic conditions than under the control conditions at high light irradiance (Fig. 3H). In contrast, the quantum yield of non-photochemical quenching at the acceptor side of PSI [Y(NA)] was lower under Mn-toxic conditions than under the control conditions at high light irradiance (Fig. 3I). These results indicated that the whole-chain photosynthetic electron transport rate was less affected under Mn-toxic conditions than under the control conditions; however, the photosynthetic electron transport reaction was limited to the donor side of PSI, and PSI was more oxidised under Mn-toxic conditions than under the control conditions.

The maximum quantum yield of PSII (Fv/Fm) was significantly, but marginally decreased under Mn-toxic conditions (Fig. 2J). The total Chl content in leaves was also decreased under Mn-toxic conditions (Fig. 2K). These results indicated that Mn toxicity stimulates PSII photoinhibition. In contrast, the total N concentration was not significantly different between the control and Mn-toxic conditions (Fig. 2L). Based on the linear function between the N and Rubisco concentrations in the leaves (Nakano *et al.,* 1997), this result indicated that the quantity of Rubisco was less affected by Mn toxicity.

### Carbohydrate metabolism in leaves grown under Mn-toxic conditions

To examine whether the suppression of photosynthesis by excessive Mn accumulation affects carbon acquisition during growth, the major carbohydrate concentrations were quantified in the leaves. Leaves were sampled at both the end of the day and the end of the night to visualise the CO_2_ assimilation activity during the day (Takagi *et al.,* 2016*a*). At the end of the day, Mn-toxicity treatments showed lower sucrose concentrations in the leaves than did the control plants (Fig. 4). In contrast, at the end of the night, the sucrose concentration was similar under both control and Mn-toxic conditions (Fig. 4). The glucose and starch concentrations were lower than the sucrose concentration under both control and Mn-toxic conditions, and significant differences were not detected between the growth conditions at the end of the day and the end of the night (Fig. 4). The sum of the sucrose, glucose, and starch concentrations was lower in leaves grown under Mn-toxic conditions at the end of the day, but the sum of these carbohydrate concentrations was similar between the growth conditions at the end of the night (Fig. 4). These results indicated that carbon acquisition is suppressed, corresponding to lower CO_2_ assimilation activities, in leaves grown under Mn-toxic conditions.

**Figure 4.**
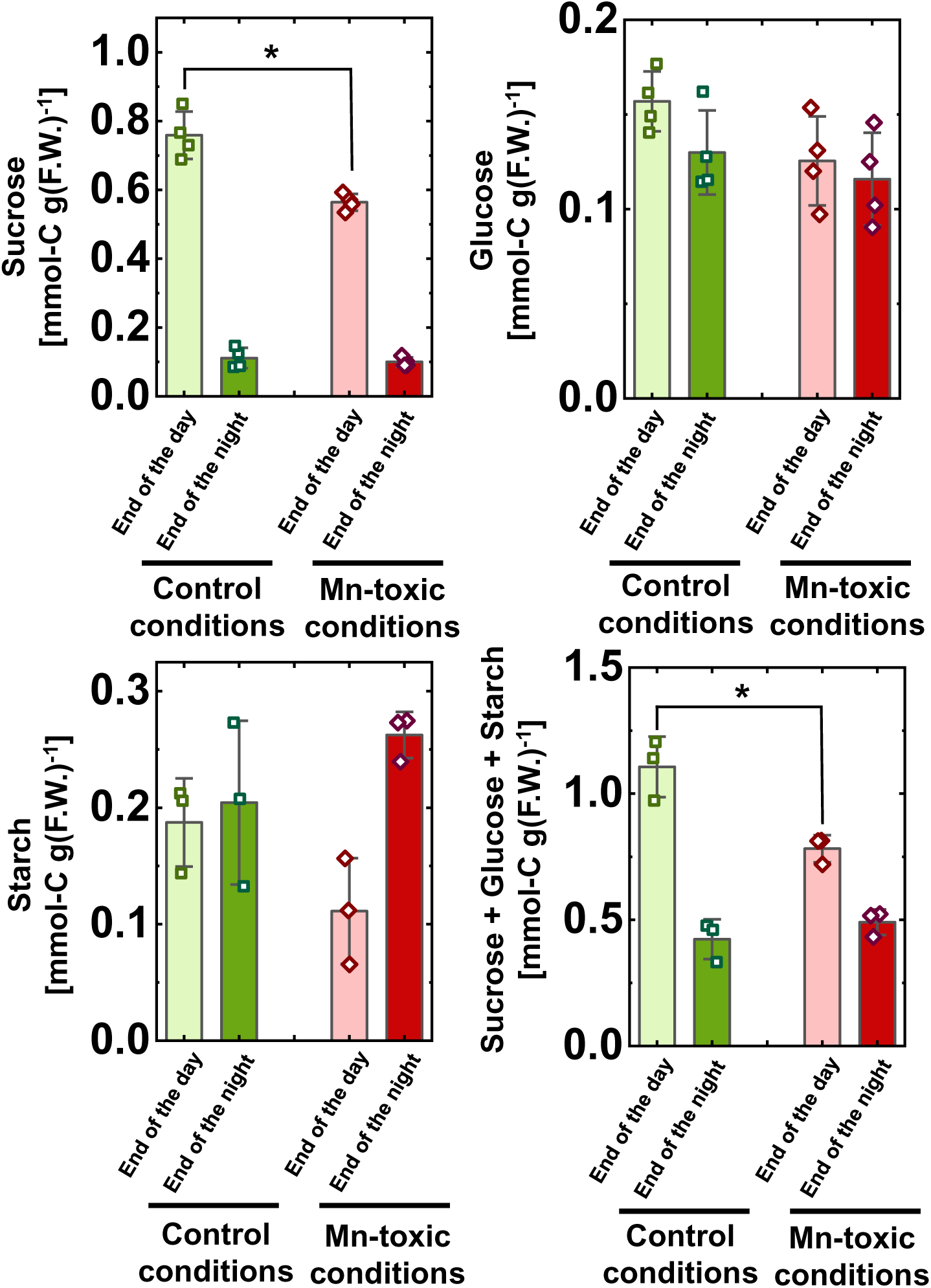
Carbohydrate content in leaf blades at the end of the day and end of the night. Data are shown as means with standard deviation (SD) (n = 3–4), and squares and diamonds show the distribution of the raw data. Asterisks show significant differences between the control and Mn-toxic conditions (*: *p* < 0.05, Kruskal–Wallis test).

### Effects of Mn toxicity on mitochondrial respiration in the leaves

The carbohydrates fixed during the day are metabolised at night and consumed at a constant rate from the beginning of the night to the dawn (Moraes *et al*., 2019). Therefore, the difference in carbohydrate concentrations between the end of the day and end of the night indicates the carbon catabolic activity during the night (Takagi *et al*., 2016*a*). To confirm whether carbon catabolism is suppressed under Mn-toxic conditions (Fig. 4), leaf respiration was examined under control and Mn-toxic conditions. The respiratory CO_2_ emission rate was decreased under Mn-toxic conditions (Fig. 5A). To examine the effect of the change in respiration activities on carbon catabolism, the leaf amino acid content was quantified during the night (Sulpice *et al*., 2014). Subsequently, the Asp, Glu, and Gly contents were significantly decreased, and the Ser content was increased under Mn-toxic conditions (Fig. 5B). These results indicated that carbon catabolism is affected by Mn-toxic conditions during the night.

**Figure 5.**
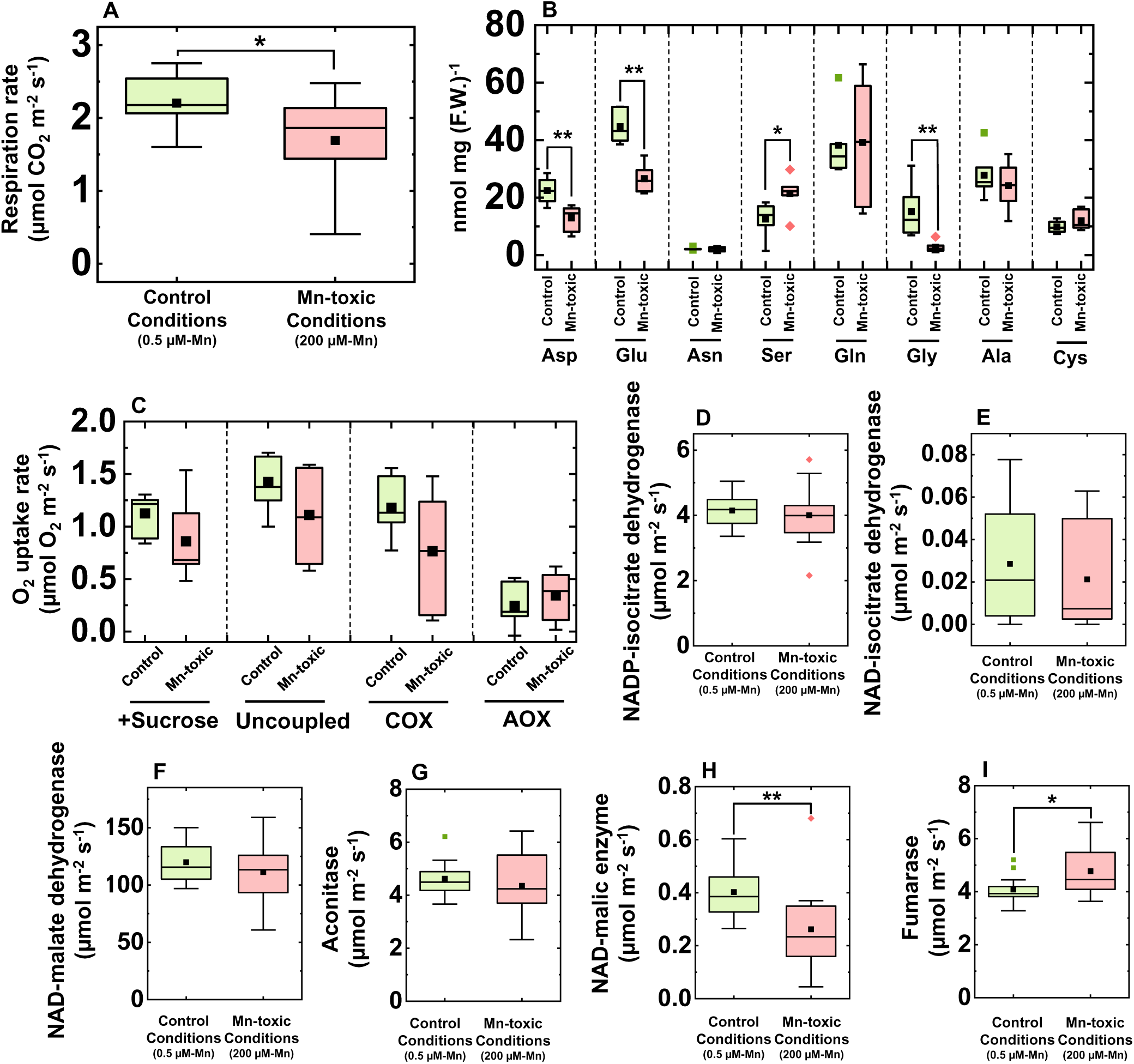
Mitochondrial respiration activities and amino acid contents in leaf blades grown under control and Mn-toxic conditions. (A) Leaf respiration rate (n = 12–13) in the dark and (B) sucrose feeding respiration activities and mitochondrial respiratory electron transport activities depending on ATP synthase (uncoupled), COX, and AOX (n = 7). (C) Amino acid content in leaves sampled in the middle of the night (n = 6). (D–I) Enzyme activities involved in the TCA cycle (n = 13–14). Data are shown as box plots, black squares indicate the mean value, and bars indicate the range of the maximum or minimum data within a 1.5× interquartile range (IQR). Green boxes indicate the results under control conditions, and red boxes indicate those under Mn-toxic conditions. Asterisks show significant differences between the control and Mn-toxic conditions (*: *p* < 0.05, **: *p* < 0.01, Kruskal–Wallis test).

Next, we investigated the effects of Mn toxicity on the mitochondrial enzymes involved in respiration. First, the respiratory electron transport activities were measured in the presence of sucrose and mitochondrial electron transport inhibitors *in vivo* using an aqueous phase O_2_-electrode (Hachiya *et al*., 2010). In the absence of inhibitors, the O_2_ consumption rate did not differ between the control and Mn-toxic conditions (Fig. 5C). CCCP dissipated mitochondrial membrane potential (Δψ_m_) and similarly stimulated O_2_ consumption rate in both leaves grown under the control and Mn-toxic conditions (Fig. 5C). The COX and AOX activities in the presence of CCCP and sucrose were determined by adding n-propyl gallate and KCN. Subsequently, the control and Mn-toxic conditions showed similar O_2_consumption rates depending on the COX and AOX activities (Fig. 5C). The effect of Mn toxicity on mitochondrial respiratory electron transport activity was reproduced when these activities were determined on a leaf fresh-weight basis (Supplementary Fig. S1). These results indicated that under sucrose feeding, the mitochondrial respiratory electron transport activities were similar between the control and Mn-toxic conditions.

To understand the effects of Mn toxicity on the enzyme activities of the TCA-cycle, the six types of TCA cycle enzyme activities were determined on both leaf area and fresh-weight basis. NADP-isocitrate dehydrogenase, NAD-isocitrate dehydrogenase, NAD-malate dehydrogenase, and aconitase showed similar activities under the control and Mn-toxic conditions (Fig. 5D-G, I, Supplementary Fig. S2). The NAD-malic enzyme activity decreased under Mn-toxic conditions (Fig. 5H, Supplementary Fig. S2), whereas the fumarase activity increased (Fig. 5I, Supplementary Fig. S2). These results indicated that Mn toxicity did not completely suppress the enzyme activities of the TCA cycle, but specifically stimulated fumarase and suppressed NAD-malic enzyme activities.

### Difference in stomatal development in the leaves between the control and Mn-toxic conditions

To examine whether the decrease in *g_s_* under Mn-toxic conditions was related to the change in stomatal structure and development, the stomatal complex on the adaxial side of a leaf was monitored. The stomatal complex, consisting of guard and subsidiary cells, was located linearly to the vein under the control conditions (Fig. 6A). However, under Mn-toxic conditions, the stomatal complex showed a more scattered distribution (Fig. 6B). The stomatal density increased under Mn-toxic conditions (Fig. 6C). In contrast, the size of the stomatal complex evaluated by the maximum major and minor axis was smaller under Mn-toxic conditions than under control conditions (Fig. 6A, B, D, E). The changes in stomatal size were consistent with a previous study evaluating rice stomata structure under excess Mn application (Lidon, 2002). These results indicated that Mn-toxic conditions change stomatal development in the leaves.

**Figure 6.**
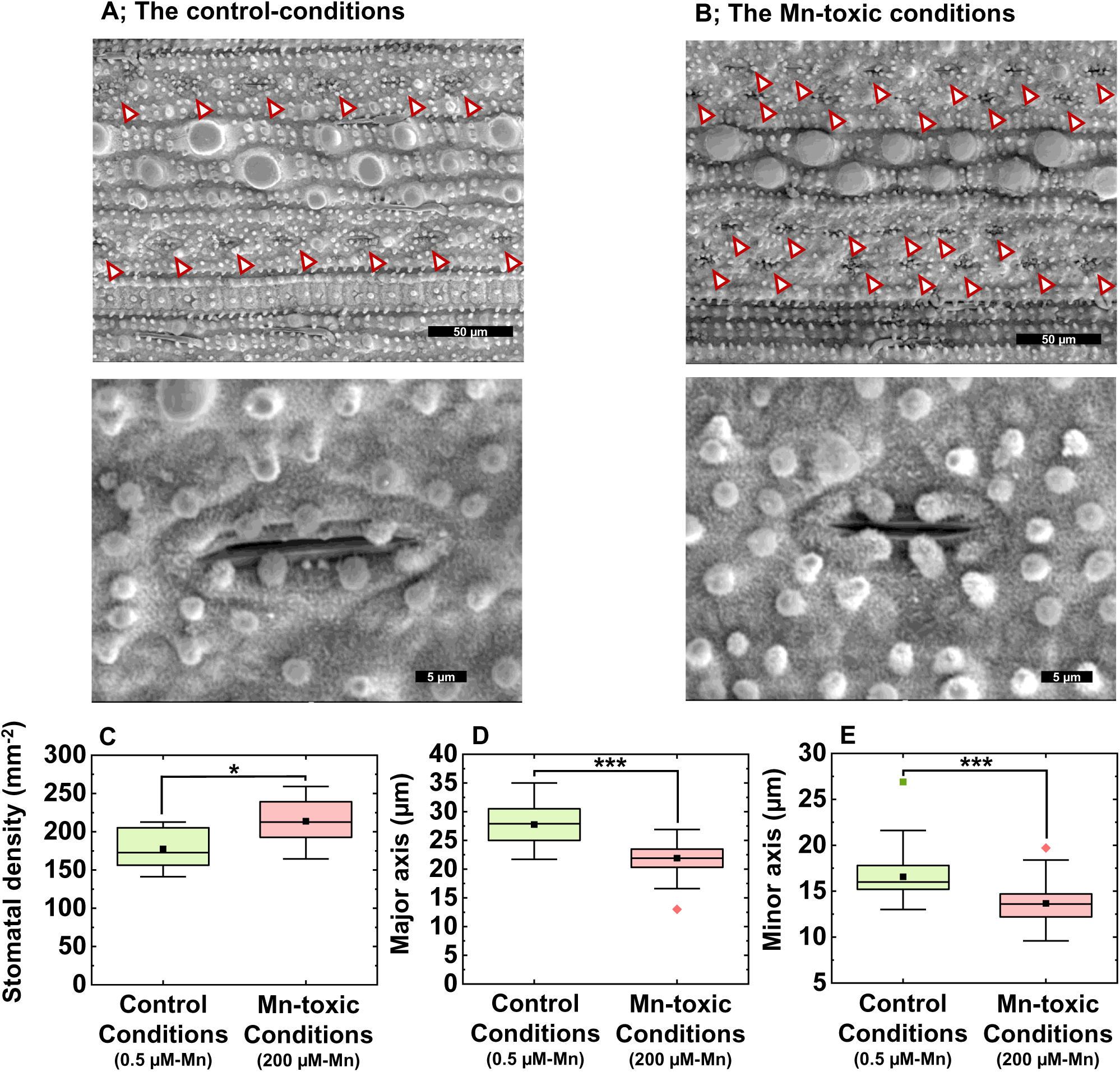
Stomatal structures and characteristics in the rice leaf blade. Pictures of the stomata on the adaxial side of the leaf blade grown under control (A) and Mn-toxic (B) conditions. The upper pictures focus on the stomatal distributions, and the lower pictures focus on the single stomata complex. (C) Showed the stomatal density (n = 10–12). (D) and (E) showed the major and the minor axis of the stomatal complex, respectively (n = 53-63). Data are shown as box plots, black squares indicate the mean value, and bars indicate the range of the maximum or minimum data within a 1.5× interquartile range (IQR). Green boxes indicate the results under control conditions, and red boxes indicate those under Mn-toxic conditions. Asterisks show significant differences between the control and Mn-toxic conditions (*: *p* < 0.05, ***: *p* < 0.001, Kruskal–Wallis test).

### Anatomical changes within a leaf blade grown under Mn-toxic conditions

Following the change in stomatal development on the leaf epidermis, the leaf anatomy was also examined. Figure 7A shows the horizontal and vertical cross-sections of the leaves against the vein. Apparent internal air spaces were observed within leaves grown under control conditions. In contrast, cells were highly condensed in leaves grown under Mn-toxic conditions, and the internal air space was hardly observed (Fig. 7A). These results showed that in addition to stomatal complex, leaf anatomical development was also altered under Mn-toxic conditions.

**Figure 7.**
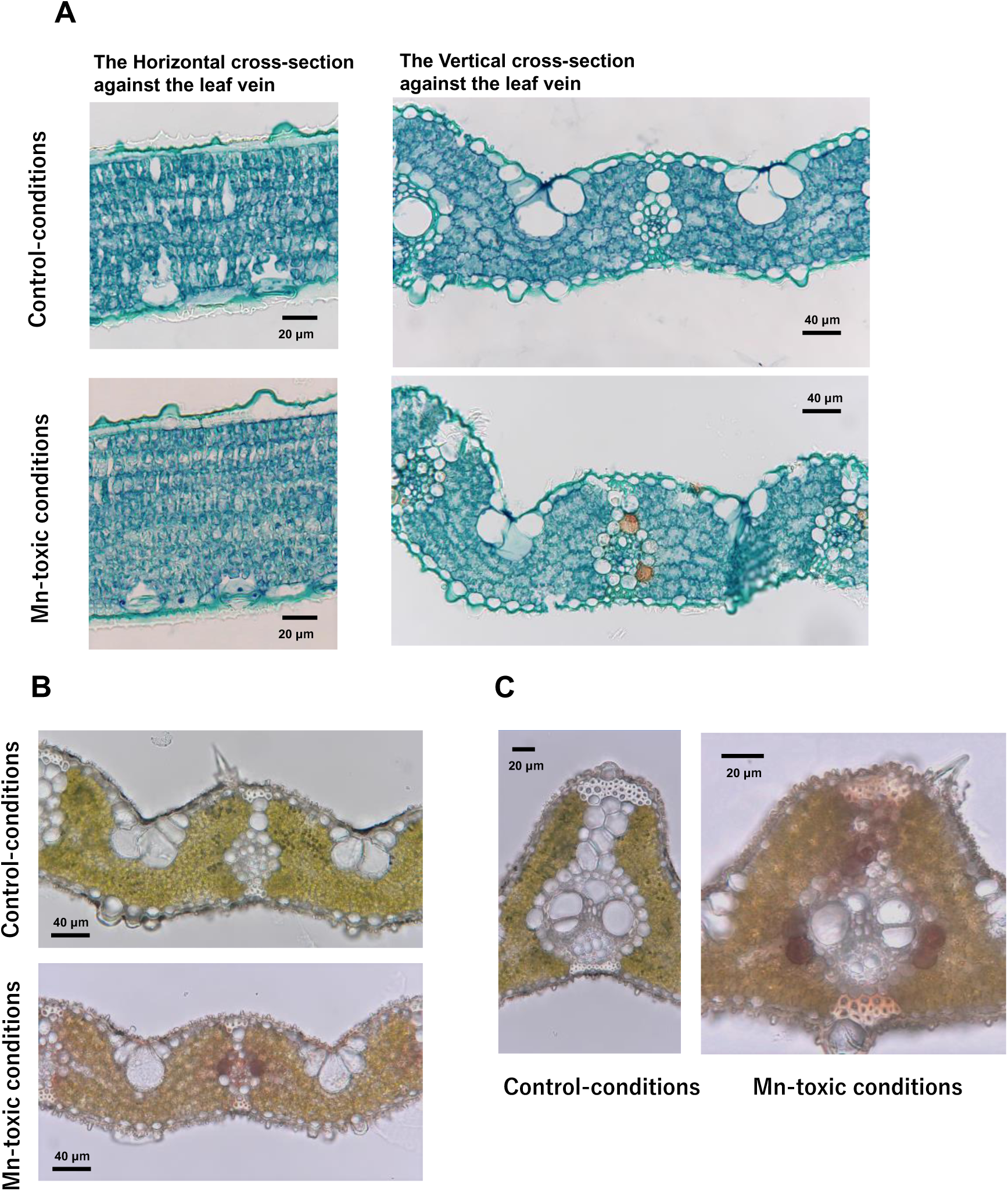
Characteristics of the leaf section grown under control and Mn-toxic conditions. (A) Fixed leaf sections sectioned from the horizontal and vertical side against the leaf vein. Fresh leaf sections of the small (B) and large (C) bundles. Black bars show the scales.

The vertical cross-section revealed that the bundle-sheath cells were brownish under Mn-toxic conditions (Fig. 7A). To investigate whether the brown spots were artifacts caused by the chemical fixing treatment, intact leaf sections were also studied. Subsequently, we found that the brownish spots natively existed in the bundle-sheath cells, both small and large bundles gown under Mn-toxic conditions (Fig. 7B, C). These results indicated that under Mn-toxic conditions, rice leaves caused apoplastic Mn toxicity-like symptoms in addition to symplastic Mn toxicity, and the symptoms are more evident in the bundle-sheath cells in rice leaves.

### Auxin content in the leaves and mRNA expression of auxin-responsive genes

Our findings demonstrated that excess Mn accumulation affects the developmental process of leaves. IAA is an important hormone that determines cell fate and differentiation in plants (Perrot-Rechenmann, 2010). Morgan et al. (1966) reported that Mn toxicity stimulates IAA oxidation activity in cotton leaves, indicating that the auxin concentration could be lowered by accelerated auxin degradation in leaves under excess Mn conditions. To examine this possibility, the IAA concentration in leaves was quantified by GC-MS. Figure 8A shows the total ion chromatograms (TIC) with fragment ion chromatograms of the 202 (*m/z*) and 319 (*m/z*) of the IAA standard and leaf extract from the control and Mn-toxic conditions (Fig 8A) (see Materials and Methods). Subsequently, this analysis revealed that the IAA concentration was significantly decreased by 76% in the leaves under Mn-toxic conditions (Fig. 8B).

**Figure 8.**
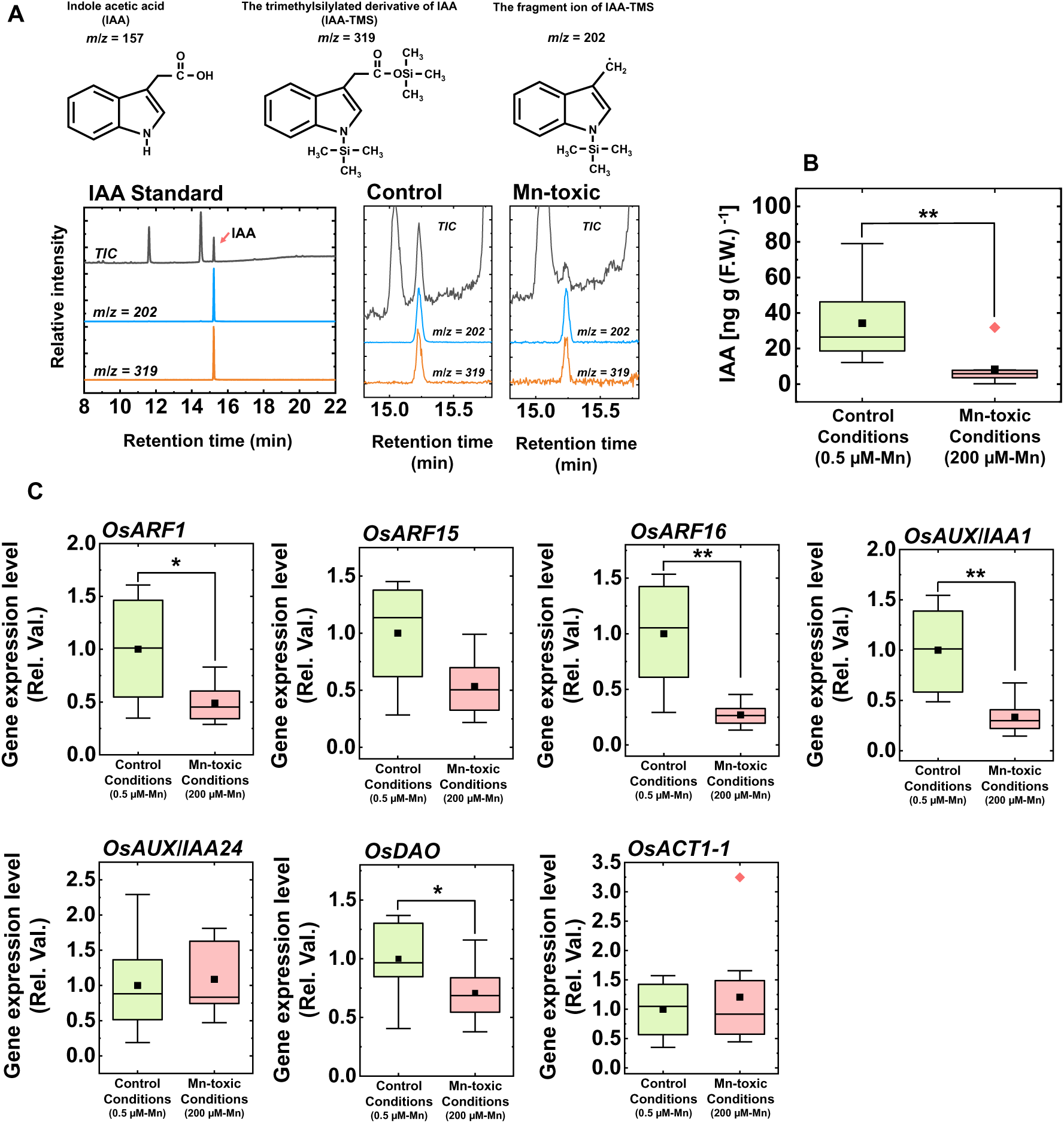
Leaf indole acetic acid (IAA) concentration and gene expression analysis relating to IAA and stomata development. (A) The result of GC-MS analysis targeting IAA, together with the chemical structures of IAA and its derivatives. The total ion chromatography (TIC) and chromatographs of *m/z* = 202 and *m/z* = 319 in the standard IAA solution and the leaf extract from plants under control and Mn-toxic conditions are shown. The retention time of the silylated IAA was 15.2 min. (B) The IAA concentrations in the leaf blade (n = 8). (D) The results of gene expression analysis involving IAA-responsive gene and stomatal patterning (n = 8). The quantified results are expressed on an *OsATC1-2* expression basis. Asterisks show significant differences between the control and Mn-toxic conditions (*: *p* < 0.05, **: *p* < 0.01, Kruskal–Wallis test).

To confirm that the decrease in IAA concentration affects physiological reactions within the leaves, the IAA-responsive gene expression was investigated. *OsARF1*, *OsARF15*, *OsARF16*, *OsAUX/IAA1,* and *OsAUX/IAA24* showed increases in their expression level with an increase in IAA concentration in the leaves (Waller *et al.,* 2002; Jain *et al.,* 2006; Wang *et al.,* 2007). Among these genes, the expression levels of *OsARF1*, *OsARF16,* and *OsAUX/IAA1* were significantly decreased under Mn-toxic conditions (Fig. 8C). *OsARF15* showed similar trends to other *ARF* genes, but the difference was not significant between the growth conditions. In contrast, *OsAUX/IAA24,* and *OsACT1-1* showed comparable expression levels between the control and Mn-toxic conditions (Fig. 8C). From these results, the auxin-responsive gene expression indicated a lower IAA concentration in leaves under Mn-toxic conditions in concert with the results of GC-MS although all genes were not responded to equally.

To regulate auxin concentration in plant cells, DIOXYGENASE FOR AUXIN OXIDATION (DAO) undertakes IAA catabolism to maintain IAA homeostasis by oxidising IAA to an inactive form, oxindole-3-acetic acid, in plant cells (Zhao *et al.,* 2013). Because Mn-toxicity stimulates oxidative IAA degradation *in vitro* (Morgan *et al.,* 1966), we hypothesised that *OsDAO* is upregulated following an increase in Mn concentration; however, the transcriptional activation of *OsDAO* was suppressed under Mn-toxic conditions (Fig. 8). The expression of *OsDAO* is activated under high IAA concentrations to maintain IAA concentrations in plant tissues (Zhao *et al.,* 2013). Based on this homeostatic response of *OsDAO*, the decrease in *OsDAO* expression under Mn-toxic conditions would be a consequence of the lower IAA concentration in leaves to avoid further inactivation of IAA.

## DISCUSSION

In acidic or waterlogged agricultural fields, Mn is easily released from the soil and causes Mn toxicity in crop plants (Fernando *et al*., 2016). In addition to the current agricultural situations, the effects of climate change, such as the rises in the atmospheric temperature and the frequency of flooding or acidic rainfall, can further accelerate Mn dissolution in soils (Fernando and Lynch, 2015). Therefore, the mechanisms of Mn toxicity should be elucidated to manipulate the future threat of increasing Mn availability on agricultural lands. To address the mechanisms of symplastic Mn toxicity, we investigated the physiological effects of high Mn application in rice. In this study, we propose that the disturbance of IAA homeostasis is one of the critical causes of symplastic Mn-toxicity in rice leaves.

### Mn toxicity disturbs IAA homeostasis and attenuates auxin signalling in the leaves

Our results showed that the IAA concentration was decreased in the leaves under Mn-toxic conditions (Fig. 8). A previous study demonstrated that IAA oxidation activity increased in leaf extracts grown under high Mn concentration conditions (Morgan *et al.,* 1966). Our observations suggest that the increase in IAA oxidation activity functions *in vivo.* In addition to the decrease in *ARF* and *AUX/IAA* gene expression levels in rice, the microRNA expression profiles revealed that miRNA160 which targets *ARF* genes was downregulated, and miRNA164 which targets the auxin-signal related F-box protein (TRANSPORT INHIBITOR RESPONSE 1, TIR1) was upregulated in the leaves of common beans under Mn toxicity (Valdés-López *et al.,* 2010). From these observations, we suggest that Mn toxicity affects the auxin signalling cascade at the transcriptional level by decreasing the IAA concentration in the leaves.

We suggest that the decrease in IAA concentration, together with the change in auxin signalling activities, causes a change in leaf anatomy and stomatal patterning/development under Mn-toxic conditions. When IAA deficiency is caused by the overexpression of GRETCHEN HAGEN3 which conjugates IAA to amino acids or the anion peroxidase which oxidatively inactivates IAA *in planta*, the leaf cell size becomes smaller and the leaf anatomy shows more condensed cell structures (Lagrimini *et al*., 1997*a*, *b*; Du *et al*., 2012). Alternatively, the transgenic plants that inhibit auxin signalling or auxin polar transport activities represent the IAA-deficient phenotype and show the abnormal leaf anatomical development (Pérez-Pérez *et al*., 2010; Guo *et al*., 2013; Qi *et al*., 2014; Muñoz-Nores *et al*., 2017). The aforementioned relationships among the IAA-deficient phenotypes and the change in leaf structure caused by attenuating auxin signal transduction are similar to those among the leaf phenotypes under Mn-toxic conditions (Figs. 3B, 6). Moreover, stomatal development is regulated by auxin signalling (Balcerowicz and Hoecker, 2014). Exogenous IAA application decreased stomatal density in a concentration-depending manner; whereas the attenuation of ARF-dependent auxin signal transduction increased it (Le *et al.,* 2014; Zhang *et al.,* 2014). Based on these results, a decrease in IAA concentrations could affect the change in leaf development and stomatal patterning/development.

Generally, the terrestrial plants with smaller stomata and higher stomata density show higher transpiration ability than those with larger stomata and lower density (Franks and Beerling, 2009). This relationship does not apply to *g_s_*and stomatal patterning (Figs. 3B, 6C); therefore, additional factors to suppress stomatal opening should exist under Mn toxicity. IAA stimulates stomatal opening by activating plasma membrane H^+^-ATPase (Jezek and Blatt, 2017), and the exogenous application of IAA stimulates stomatal opening in the leaves (Klein *et al*., 2003). Therefore, the decrease in *g_s_* could be caused by reducing the IAA concentration in the leaves. However, the regulation of stomatal function is quite complex, and the ROS and organic acid composition are also determinants of stomatal movement (Jezek and Blatt, 2017). Considering the stimulation of oxidative stress under Mn toxicity (see the following discussion), the decrease in IAA would not be the only cause of stomatal dysfunction. Interestingly, fumarase activity increased under Mn toxicity (Fig. 5I). Because the increase in malate/fumarate ratio is important for stomatal opening (Araújo *et al*., 2011), the increased fumarase activity might be a counteraction toward stomatal dysfunction under Mn toxicity.

Currently, we cannot determine how the IAA concentration is lowered under Mn toxicity at the molecular level. Further research is required on a molecular genetic basis to elucidate the key factors that cause IAA deficiency. We now consider the possibility that IAA oxidation can be undertaken by unknown proteins such as class III PODs as reported previously (Ray, 1958; Lagrimini *et al*., 1997*a*, *b*; Tognetti *et al*., 2012). Because IAA is transported through the apoplast, the increase in POD activity in the apoplast might influence IAA catabolism under Mn toxicity (Robert and Friml, 2009).

### Suppression of photosynthesis is caused by stomatal dysfunction in rice

We suggest that the simultaneous prolonged limitation of the CO_2_ assimilation reaction and stimulation of photorespiration caused oxidative stress by ROS under Mn toxicity. Under Mn toxicity, ROS-scavenging enzyme activities are upregulated in various terrestrial plants (Gonzáletz *et al*., 1988; Lei *et al*., 2007; Santos *et al*., 2017). This response indicates that oxidative stress is stimulated by ROS in cells under Mn toxicity. Furthermore, Houtz *et al*. (1988) showed an increase in the RuBP concentration in leaves under Mn toxicity. This result indicated that the turnover of photosynthates is limited by Rubisco activity (Suzuki *et al*., 2012). Here, *g_s_* was lowered and the CO_2_ assimilation rate was limited under Mn-toxic conditions (Fig. 2B). In contrast to the CO_2_ assimilation rate, which showed a 39% decrease at the highest light irradiance, Y(II) only decreased by 9% at the same light intensity (Fig. 3). This difference would have been caused by the difference in the electron distribution between carboxylation and oxygenase reaction by Rubisco (Takagi *et al*., 2016*b*; Wada *et al*., 2019). Even when the CO_2_ availability within chloroplasts is limited by stomatal closure, O_2_ can diffuse within the leaves; therefore, Rubisco drives photorespiration (Zivcak *et al*., 2013). Under such conditions, and the *proton motive force* is built up to stimulate luminal acidification of the thylakoid membranes owing to the lower ATP consumption rate in the photorespiration reaction compared with that in the carboxylation reactions in the Calvin-Benson cycle (Takagi *et al*., 2016*b*, 2017). Luminal acidification is represented by an increase in Y(ND) (Fig. 3), which is increased by limiting the electron flow from plastoquinone to cytochrome *b_6_f* depending on the luminal acidification (Kramer *et al*., 1999), under Mn-toxic conditions. These results indicated that Mn toxicity stimulates photorespiration via stomatal closure. The increase in photorespiration activity can also be supported by the change in the Gly/Ser ratio under Mn-toxic conditions (Fig. 5B). Wada *et al*. (2019) reported that the limitation of the CO_2_ assimilation rate by drought stress with a decrease in *g_s_* stimulates PSI photoinhibition rather than PSII photoinhibition by ROS. A similar situation would occur under Mn toxicity, which is why the decrease in PSII activity is modest, although PSI inhibition is accentuated under Mn-toxic conditions (Kitao *et al*., 1997; Lidon *et al*., 2004; Millaleo *et al*., 2013). Previous studies have shown that Mn toxicity targets PSI by attenuating Fe absorption (Andressen *et al*., 2018). However, the concentration was maintained within a sufficient range for Fe nutrition in rice [from 70 to 300 µg g^-1^ (D.W.)] (Fageria and Stone, 2008). Furthermore, a decrease in the PSI content caused an increase in Y(NA), instead of Y(ND) during steady-state photosynthesis (Brestic *et al*., 2015). Hence, we suggest that the decrease in PSI content due to Mn toxicity would be caused not only by the Fe status but also by PSI photoinhibition, as a consequence of limited CO_2_ assimilation. Other nutrients such as Ca, Mg and Zn were also reported to be deficient under Mn toxicity (Broadley *et al*., 2012). We did not observe a decrease in these minerals in the leaves, but rather increased Mg, Cu, and Zn concentrations under Mn toxicity (Fig. 2). Because Cu/Zn-type SOD activities are upregulated under Mn toxicity (Gonzáletz *et al*., 1988; Lei *et al*., 2007; Santos *et al*., 2017) and Mg supply rescues Mn toxicity symptoms (Broadley *et al*., 2012), the requirement of these minerals might be upregulated.

Our results suggest that chloroplasts have homeostatic systems to evade excessive Mn accumulation. *In vitro* studies have shown that excess free Mn stimulates ROS production by PSII and causes severe oxidative damage to thylakoid lipids and PSII (Panda *et al.,* 1987). Although the Mn concentration was high (Fig. 2), clear evidence of severe damage to PSII was not detected here (Fig. 3J). Furthermore, Mn is to bind Rubisco, and an *in vitro* study indicated that the substitution of Mg^2+^ with Mn^2+^ decreases the specificity of Rubisco for CO_2_ (Jordan and Ogren, 1983; Bloom and Kameritsch, 2017). Previous studies have suggested that Mn binding to Rubisco causes a decrease in photosynthesis under Mn toxicity (Houtz *et al.,* 1988; Kitao *et al.,* 1997). However, the contribution of Mn-binding Rubisco to decrease CO_2_ assimilation reactions remains unknown under Mn-toxic conditions *in vivo* (Houtz *et al.,* 1988; Chatterjee *et al.,* 1994). To address this question, we simulated the change in CO_2_ fixation rate between Mg^2+^- and Mn^2+^-binding Rubisco using photosynthetic biochemical model (von Caemmerer and Farquhar, 1981). Calculated CO_2_ assimilation rate using Mn^2+^-binding Rubisco kinetics was negative value below Cc = 50 Pa and greatly decreased compared to calculated CO_2_ assimilation rate using Mg^2+^-binding Rubisco kinetics (Supplementary Fig. S3). However, such a drastic decrease in CO_2_ assimilation rate was not observed (Fig. 3). Although Mn^2+^-binding Rubisco showed a higher photorespiration rate than Mg^2+^-binding Rubisco, this difference cannot compensate the difference in CO_2_ assimilation rate. Moreover, investigation of the relationship between *g*_s_ and the CO_2_ assimilation rate did not show a decrease in the CO_2_ assimilation rate independent of the change in *g_s_* (Supplementary Fig. S4). These considerations reconciled the cause of the decrease in CO_2_ assimilation as decrease in *g_s_*, and Rubisco is protected from excess Mn in chloroplasts. To protect chloroplasts from excess free Mn accumulation, the homeostatic function of Mn^2+^ in the chloroplasts should function. At this point, two mechanisms can be considered. The first mechanism is binding Mn to the thylakoid membranes. Although this modulates membrane structure of the thylakoid membranes, no inhibitory effect on photosynthetic activity was observed (Lidon and Teixeira, 2000*a*, *b*). The binding of Mn to the thylakoid membrane could be a storage system for Mn homeostasis. The second mechanism is the sequestration or limiting Mn transport within chloroplasts. Indeed, Mn distributed to the chloroplasts was decreased under Mn-toxic conditions in rice (Führs *et al.,* 2010). These homeostatic systems would prevent the deleterious effects of Mn^2+^ within the chloroplasts.

### Carbon catabolism due to reduced carbohydrates suppresses growth

A decrease in respiration rate under Mn toxicity has been reported in wheat and cotton plants (Sirkar and Amin, 1974; Macfie and Taylor, 1992). We also observed a decrease in leaf respiration, but not in sucrose feeding (Fig. 5). Moreover, no decreases in mitochondrial enzyme activities, including TCA cycle and respiratory electron transport chain enzymes, were observed, except for the NAD-malic enzyme (Fig. 5). The physiological importance of the NAD-alic enzyme is highly limited in C3 plants, and a pronounced phenotype has not been observed in NAD-malic enzyme mutants in *Arabidopsis* (Tronconi *et al*., 2008). This could be because of the lower contribution to the supply of pyruvate into the TCA cycle by NAD-malic enzyme activity (Williams *et al.,* 2008). These observations suggested that the significant decrease in NAD-malic enzyme activity could be independent of the growth defect and decreased respiration rate under Mn-toxic conditions (Figs. 1, 5). However, the change in amino acid composition could be influenced by a decrease in the NAD-malic enzyme (Tronconi *et al.,* 2008). From these observations, the decreased respiration rate was caused by a decrease in carbohydrate concentrations due to the suppression of photosynthesis (Sulpice *et al.,* 2014). Carbohydrate catabolism during the night is a determinant of plant growth (Takagi *et al.,* 2016*a*). We suggest that a decrease in the capacity of carbon catabolism causes growth defects in rice under Mn toxicity. Interestingly, Hibberd and Quick (2002) reported that similar to C4 plants, the NAD-malic enzyme activity is high in the bundle-sheath cells in C3 plants. Here, we found that the bundle-sheath cells turned brown under Mn-toxic conditions (Fig. 7B, C). Based on a previous study on apoplastic Mn toxicity, brown spots contained oxidised phenol, oxidised Mn, and callose. Fernando *et al*. (2016) showed that wheat leaves concentrate Mn in the bundle-sheath cell. However, this specific localisation was not observed in soybean and canola, suggesting that the bundle-sheath cell could function as storage organs for attenuating Mn toxicity in monocots. The Mn accumulation in the bundle sheath cell could cause the suppression of the NAD-malic enzyme activity in leaves. These results provide insight for elucidating the function of bundle-sheath cells in C3 monocots.

## CONCLUDING REMARKS

We proposed that the disturbance of IAA homeostasis is an initial cause of the inhibition of CO_2_ assimilation and photoinhibition caused chlorosis in the leaves (Fig. 9). As a short-term effect, the decrease in IAA suppresses stomatal opening. Subsequently, the decrease in IAA concentration modifies leaf anatomy and stomatal development in newly emerged leaves as a long-term effect. Because leaf anatomy determines the CO_2_ pathway for efficient CO_2_ absorption by the chloroplasts, the lower internal air space could further lower the CO_2_ fixation efficiency under Mn toxicity (Earles *et al.,* 2018). The decreased CO_2_ assimilation rate and increased photorespiration rate decelerate carbon acquisition for catabolism; thus, vegetative growth is suppressed. To our knowledge, there are limited studies focusing on the relationship between auxins and Mn toxicity (Morgan *et al.,* 1966). Tsunemitsu *et al*. (2018) reported that the *osmtp11* mutant decreased grain fertility and yield in rice. *OsMTP11* undertakes Mn sequestration to the endoplasmic reticulum and is highly expressed in inflorescences, including the anthers, pistil, lemmas, paleas, and ovaries (Tsunemitsu *et al.,* 2018). Together with our conclusion, Mn toxicity may also affect grain fertilisation by disturbing IAA homeostasis. Not only the photosynthetic measurements but also the developmental analyses from leaf emergence to the reproductive phase would provide more insights into the mechanisms of the symplastic Mn-toxicity by focusing on IAA.

**Figure 9.**
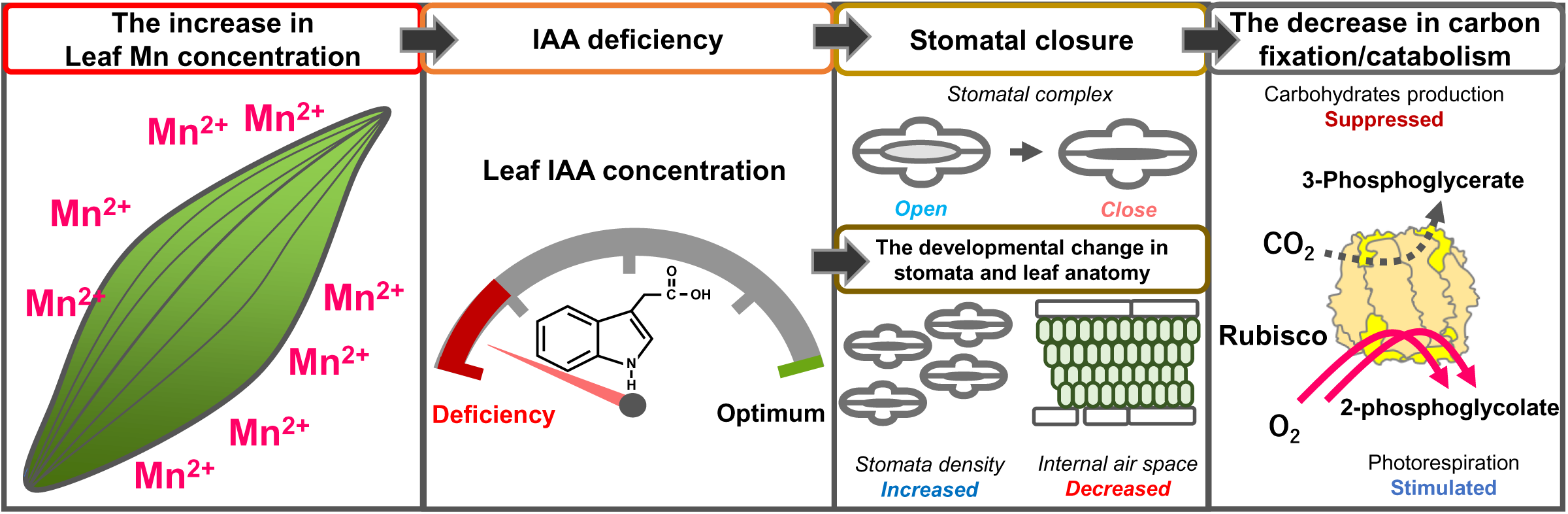
Visual scheme of symplastic Mn toxicity to suppress CO_2_ assimilation. When Mn is excessively accumulated in leaves, the leaf IAA concentration is lowered, which might cause the stimulation of IAA degradation activities under Mn toxicity (Morgan *et al.,* 1996). As a short-term effect, the decrease in IAA concentration affects stomatal opening (Blatt and Thiel, 1994; Klein *et al.,* 2003). Subsequently, as a long-term effect, the auxin-dependent signal transduction involving ARF transcriptional factor is perturbated in the leaves to cause an IAA-deficient phenotype, changing both stomatal and leaf anatomical structures (Le *et al*., 2014; Zhang *et al.,* 2014). The change in stomatal function and leaf structure severely limits CO_2_ diffusion to the chloroplasts, and CO_2_ assimilation by Rubisco is inhibited. In contrast, the photorespiration reaction limits the photosynthetic electron activities to cause ROS production (Wada *et al.,* 2019). The decreased sugar production efficiency in photosynthesis suppresses sugar catabolism. Therefore, growth is inhibited under Mn-toxic conditions.

## SUPPLEMENTARY DATA

**Figure S1;** The sucrose-feeding respiration activities and mitochondrial respiratory electron transport activities depending on ATP synthase (uncoupled), COX, and AOX based on leaf fresh weight

**Figure S2;** The enzyme activities involved in the TCA cycle calculated on the fresh weight basis.

**Figure S3;** The simulated CO_2_ assimilation in leaves containing Mg-binding Rubisco and Mn-binding Rubisco.

**Figure S4;** The relationship between *g_s_* and CO_2_ fixation rate under the control and the Mn-toxic conditions.

**Table S1;** The primer list for the gene expression analysis in rice leaf blade

**Table S2;** Mg^2+^-binding Rubisco and Mn^2+^-binding Rubisco kinetics.

## ACKNOWLEDGEMENTS

The authors thank Editage (https://www.editage.jp) for English language editing of the manuscript. The authors also thank Prof. Yukio Ishikawa, Prof. Yutaka Okumoto, Prof. Takeo Yamakawa, and Associate Prof. Shuji Sano at Setsunan University for their kind technical support of this study. This work was supported by the Japan Society for the Promotion of Science (JSPS) research fellowship (JSPS KAKENHI Grant No. 18J00852 [DT] and KAKENHI Grant No.16H06379 [AM]).

## Author contributions

Conceptualisation: DT; investigation: DT, KI, MS, TU, TF, YT, MK, and KO; original draft: DT; writing, review, and editing: DT, KI, MS, TU, TF, YT, MK, KO and AM; funding acquisition: DT and AM.

## Data availability statements

All data supporting the findings of this study are available within the article and its supplementary figures and tables published online.

## ABBREVIATIONS

AOX: alternative oxidase
ARF: auxin response factor
AUX/IAA: auxin/indoleacetic acid
Cc: chloroplastic CO_2_ partial pressure
COX: cytochrome *c* oxidase
Ci: internal CO_2_ concentration in leaves
DAO: dioxygenase for auxin oxidation
D.W.: dry weight
F.W.: fresh weight
Fv/Fm: maximum quantum yield of photosystem II
GC-MS: gas chromatography-mass spectrometry
*g_s_*: stomatal conductance
IAA: indole acetic acid
ICP-OES: inductively coupled plasma optical emission spectroscopy
HPLC: high-performance liquid chromatography
PODs: class III peroxidases
PS: photosystem
ROS: reactive oxygen species
SEM: scanning electron microscope
SOD: superoxide dismutase
TCA: tricarboxylic acid
Y(II): quantum yield of photosystem II
Y(I): quantum yield of photosystem I
Y(NPQ): quantum yield of non-photochemical quenching
Y(NO): quantum yield of non-radiative decay
Y(ND): quantum yield of non-photochemical quenching at the donor-side of photosystem I
Y(NA): quantum yield of non-photochemical quenching at the acceptor-side of photosystem I

