## Supplemental Tables and Figures for "Manganese toxicity disrupts indole acetic acid homeostasis and suppresses CO_2_ assimilation reaction in rice plants"

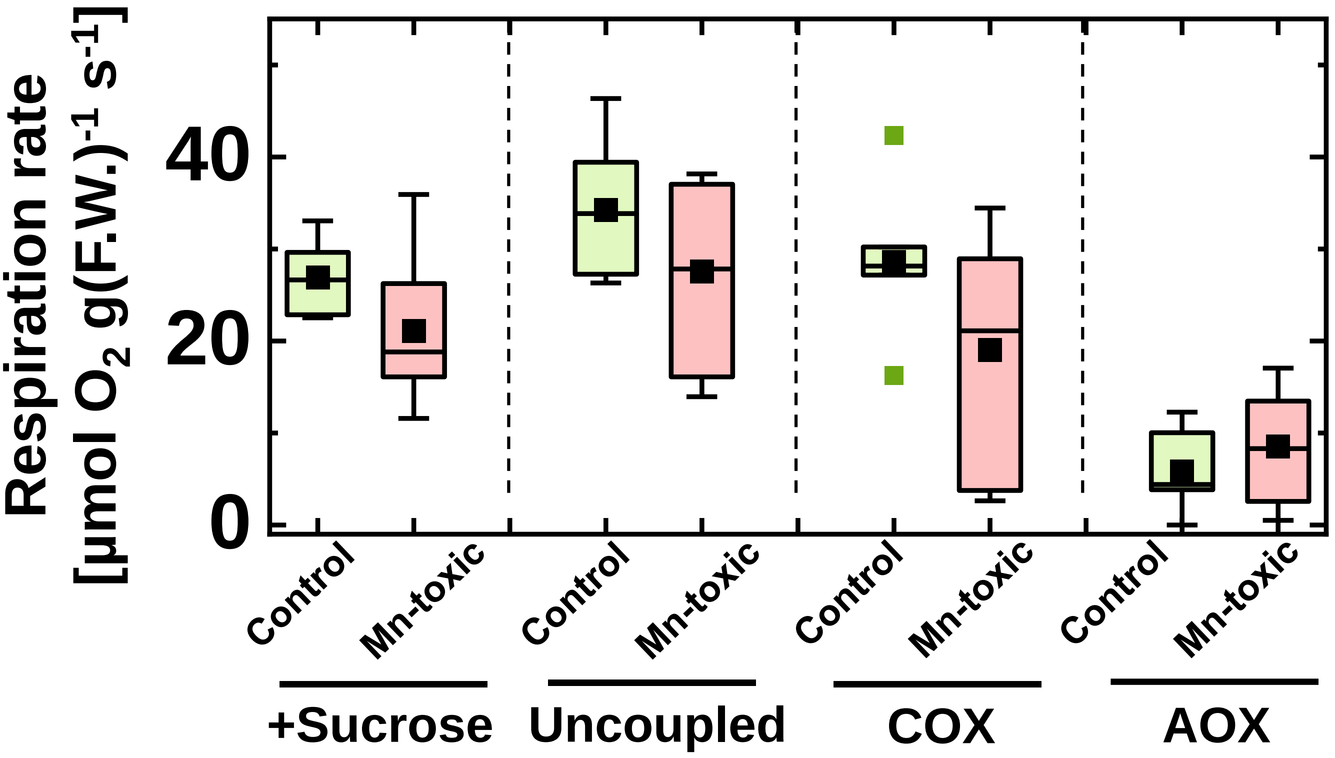


**Figure S1**

The sucrose-feeding respiration activities and mitochondrial respiratory electron transport activities depending on ATP synthase (uncoupled), COX, and AOX based on leaf fresh weight (n = 7). Data are shown as box plots, and black squares indicate the mean value, and bars indicate the 1.5 × IQR (interquartile range) of the data. Green boxes indicate the results of the control-conditions, and red boxes indicate those of the Mn-toxic conditions.

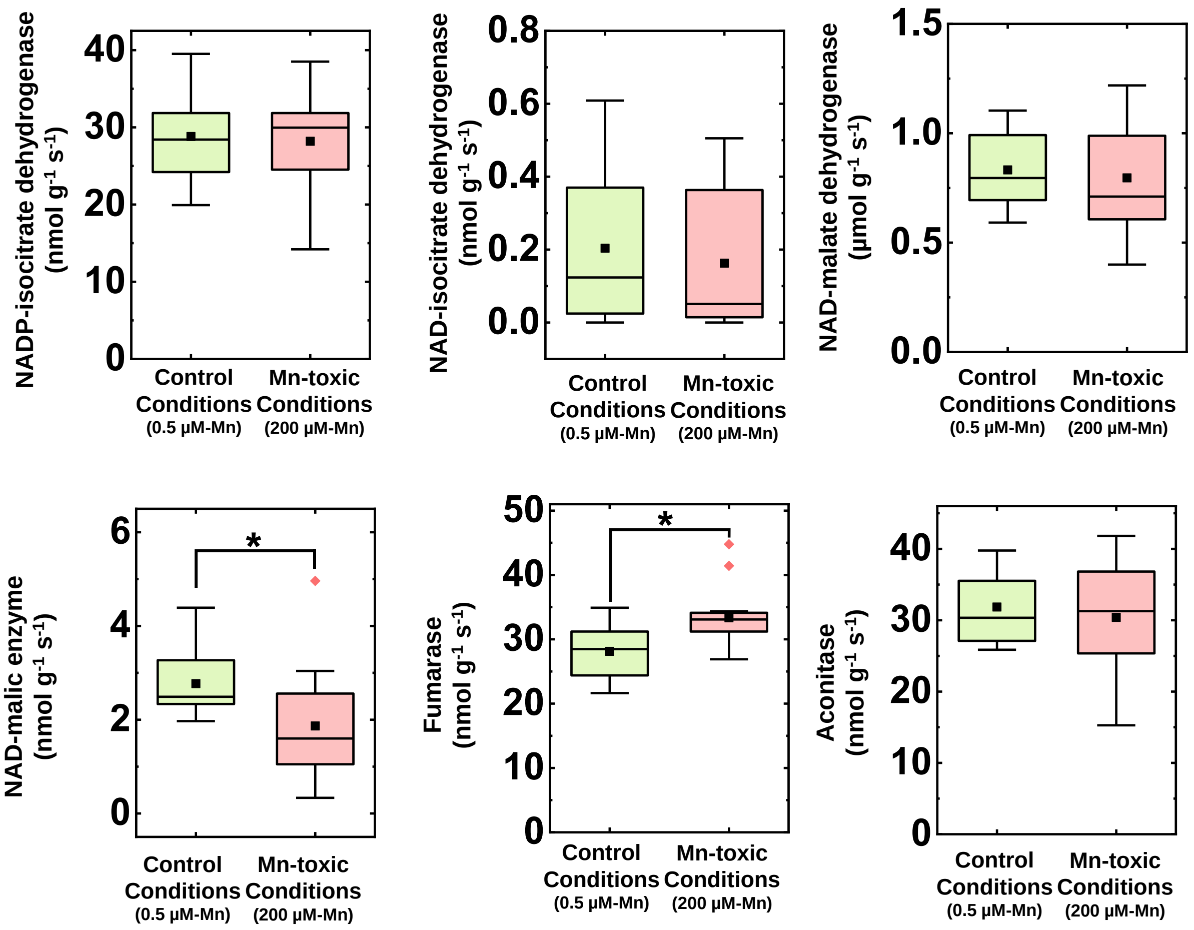


**Figure S2**

The enzyme activities involved in the TCA cycle, respectively (n = 13-14). The activities were expressed on a leaf-fresh weight basis. Data are shown as box plots, and black squares indicate the mean value, and bars indicate the 1.5 × IQR (interquartile range) of the data. Green boxes indicate the results of the control-conditions, and red boxes indicate those of the Mn-toxic conditions. Asterisks showed significant differences between the condition- and the Mn-toxic conditions (*; $p$ < 0.05, **; $p$ < 0.01, Kruskal-Wallis test).

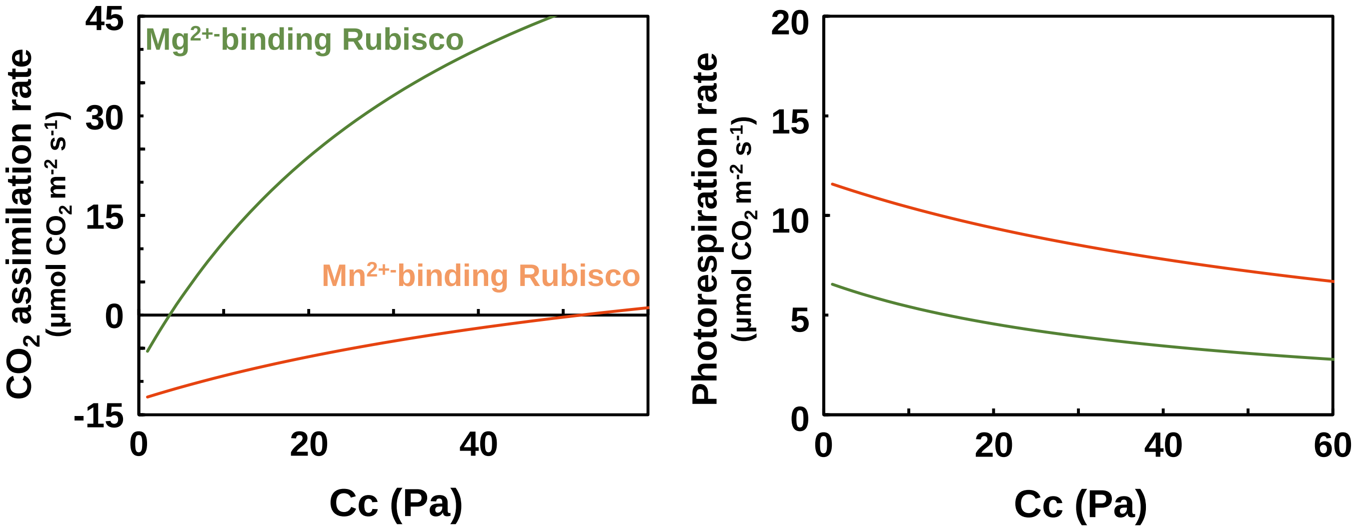


**Figure S3**

The simulated Rubisco-limited $CO_2$ assimilation rate and photorespiration rate in leaves containing $Mg^{2+}$ binding Rubisco and $Mn^{2+}$-binding Rubisco. Rubisco-limited $CO_2$ assimilation rate were calculated from Equation (1). We assumed that Rubisco content was the same as in our previous study (Suganami et al. 2021). $Mg^{2+}$/$Mn^{2+}$ binding Rubisco kinetics were summarized in Table. S2. Photorespiration rate were calculated from Equation (3). The green line indicates the simulation using $Mg^{2+}$-binding Rubisco kinetics and the pink line indicates the simulation using $Mn^{2+}$-binding Rubisco kinetics.

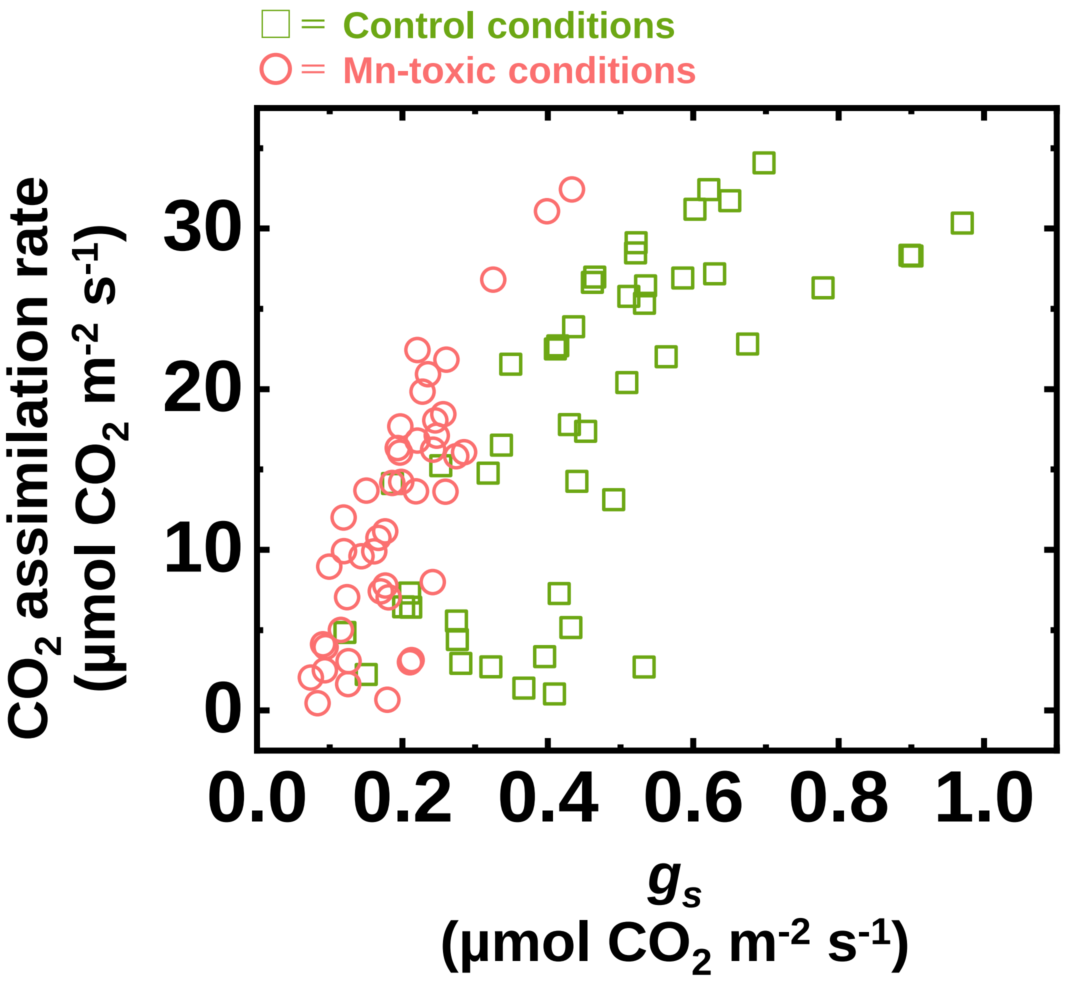


**Figure S4**

The relationship between $g_s$ and $CO_2$ assimilation rate under the control and the Mn-toxic conditions. The data shown here is identical to the results shown in Figure 3. Green squares indicate the result of the control conditions, and pink circles indicate the result of the Mn-toxic conditions.

**Supplemental Table S1**

The primer list for the gene expression analysis in rice leaf blade

| Gene name | Gene ID | | Primer sequence (5' to 3') |
|---|---|---|---|
| *OsARF1* | Os11g0523800 | Left | ACTGGATATGAGCCGTCAGC |
| | | Right | GGTGTCTTCGTGGTTGACCT |
| *OsARF15* | Os05g0563400 | Left | TCCCTCCACTGGATTACAGC |
| | | Right | CAAATGCACTCCAACCTGTG |
| *OsARF16* | Os01g0236300 | Left | TGACCCGGATCAAGAGAATC |
| | | Right | GGATCTTGCAGAAGGAGTGC |
| *OsAUX/IAA1* | Os01g0178500 | Left | CGCTCCAGGACAAGTTCTTC |
| | | Right | GTACTCCGTCCCGTTCACC |
| *OsAUX/IAA24* | Os07g0182400 | Left | AAGGCACAGGTGGTAGGATG |
| | | Right | ATCACCACCCTTCTTGGTTG |
| *OsDAO* | Os04g0475600 | Left | GAGAGGATGCACTCGCTGAT |
| | | Right | ACGGAGTCCTGCGTGTAGTT |
| *OsACT1-1* | Os03g0718100 | Left | ATAGCATGGGGGAGAGCATA |
| | | Right | CGTCTGCGATAATGGAACTG |
| *OsACT1-2* | Os03g0718100 | Left | TCCATCTTGGCATCTCTCAG |
| | | Right | GTACCCGCATCAGGCATCTG |

**Table S2** $Mg^{2+}$-binding Rubisco and $Mn^{2+}$-binding Rubisco kinetics.

| | **$Mg^{2+}$-binding Rubisco** | **$Mg^{2+}$-binding Rubisco** |
|---|---|---|
| **Vc (mol mol$^{-1}$ Rubisco s$^{-1}$)** | 17.3 | 3.9 |
| **Vo (mol mol$^{-1}$ Rubisco s$^{-1}$)** | 5.7 | 5.3 |
| **Kc (Pa)** | 23.6 | 13.4 |
| **Ko (kPa)** | 26.2 | 4.3 |

$Mg^{2+}$-binding Rubisco kinetics were taken from Makino et al. (1988). $Mn^{2+}$-binding Rubisco kinetics were calculated by multiplying the average ratio of $Mn^{2+}$/$Mg^{2+}$-binding Rubisco kinetics and $Mg^{2+}$-binding Rubisco kinetics. The average ratio of $Mg^{2+}$/$Mn^{2+}$-binding Rubisco kinetics were taken from Bloom and Kameritsch (2017).
